# An Anatomical and Physiological Basis for Flexible Coincidence Detection in the Auditory System

**DOI:** 10.1101/2024.02.29.582808

**Authors:** Lauren J Kreeger, Suraj Honnuraiah, Sydney Maeker, Siobhan Shea, Gord Fishell, Lisa V Goodrich

**Affiliations:** Harvard Medical School, Department of Neurobiology, Boston, MA 02115, USA; Stanley Center for Psychiatric Research, Broad Institute of MIT and Harvard, Cambridge, MA 02142, USA

## Abstract

Animals navigate the auditory world by recognizing complex sounds, from the rustle of a predator to the call of a potential mate. This ability depends in part on the octopus cells of the auditory brainstem, which respond to multiple frequencies that change over time, as occurs in natural stimuli. Unlike the average neuron, which integrates inputs over time on the order of tens of milliseconds, octopus cells must detect momentary coincidence of excitatory inputs from the cochlea during an ongoing sound on both the millisecond and submillisecond time scale. Here, we show that octopus cells receive inhibitory inputs on their dendrites that enhance opportunities for coincidence detection in the cell body, thereby allowing for responses both to rapid onsets at the beginning of a sound and to frequency modulations during the sound. This mechanism is crucial for the fundamental process of integrating the synchronized frequencies of natural auditory signals over time.

## Introduction

Perception depends on the ability of neurons to encode discrete features of the sensory environment. To generate accurate percepts of complex auditory stimuli, neurons must compute both what frequencies are present and when those frequencies occur across multiple time scales. For instance, overlapping sounds, such as two competing speakers in a noisy room, are distinguished perceptually by correctly binding frequencies with coherent onsets and synchronized temporal modulations^1,2^. Such computations require coincidence detection that can encode co-occurring frequencies with submillisecond precision. Frequency information is communicated by spiral ganglion neurons (SGNs), whose central axons, also called auditory nerve fibers, project through the eighth nerve, bifurcate, and spread tonotopically to fill each division of the cochlear nucleus complex (CNC)^3–7^ (**Fig. 1A**). In addition, SGNs fall into physiologically distinct subtypes that are recruited at different intensities, allowing sounds to be detected across a wide dynamic range ^8–14^. Many target neurons receive SGN inputs from a limited range of frequencies, encoding only a fraction of the frequency components within a stimulus. This presents a challenge for auditory circuits which must ultimately bind co-occurring frequencies while retaining sequence information in order to locate and recognize sounds. Understanding how these first computations are made is a key step towards deciphering the basis of perception.

**Figure 1.**
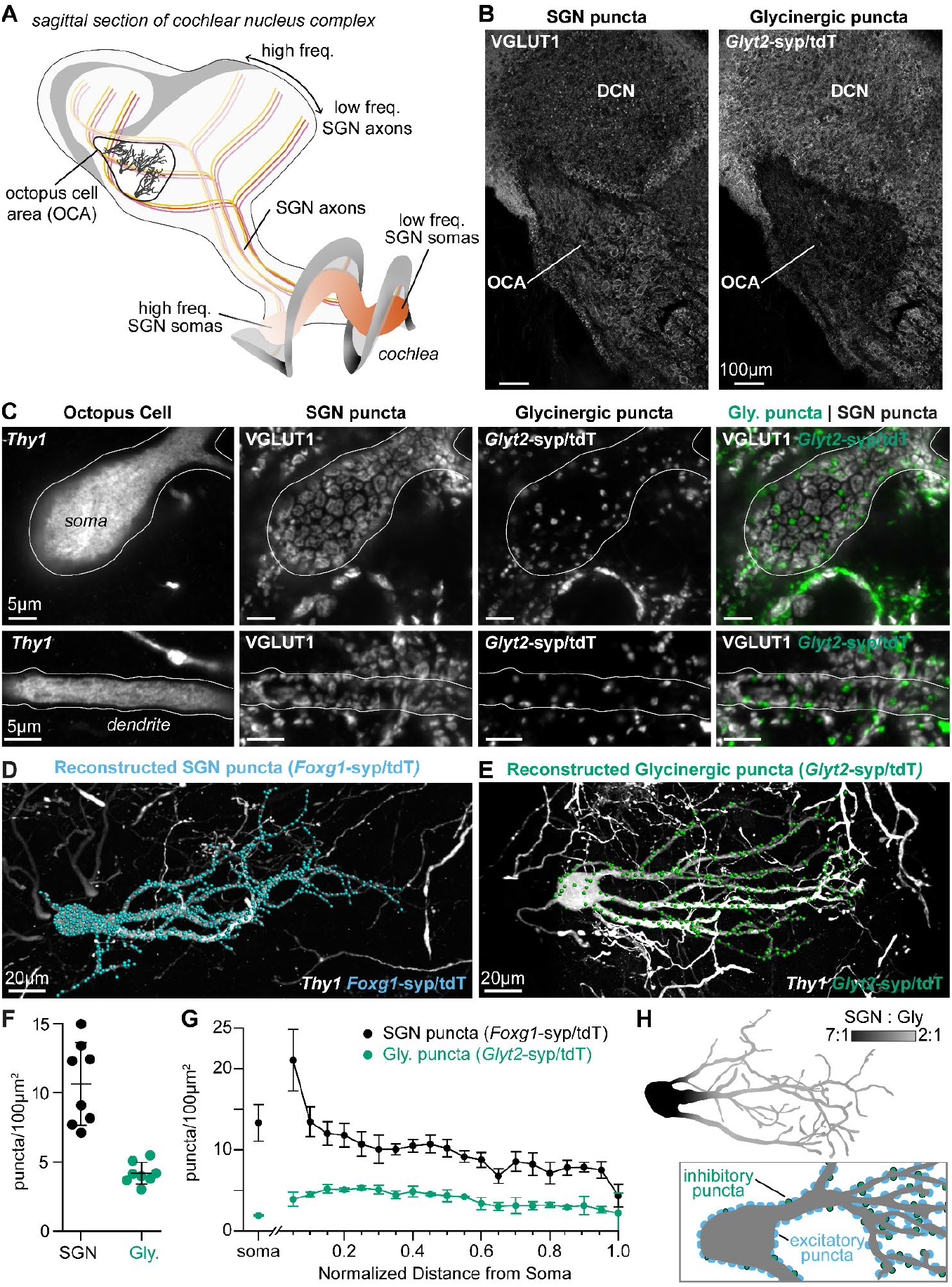
Excitatory and inhibitory synapses to octopus cells form two domains. **(A)** Illustration of spiral ganglion neuron (SGN) central axons branching within a parasagittal section of the mouse cochlear nucleus complex (CNC). SGN somas in the cochlea are tonotopically organized according to frequency. Axons remain organized throughout the ventral (VCN) and dorsal (DCN) divisions of the CNC. Octopus cells (inset) are found in the octopus cell area (OCA) of the VCN. **(B)** Excitatory SGN puncta labeled with a VGLUT1 antibody (left) and glycinergic puncta labeled with *Glyt2*^Cre^-dependent syp/tdT (*Glyt2*-syp/tdT; right) in a parasagittal section of the CNC. The teardrop-shaped OCA is not devoid of inhibitory inputs, although they are less prominent than in the surrounding CNC. **(C)** A *Thy1* sparsely labeled octopus cell with excitatory SGN (VGLUT1) and inhibitory (*Glyt2*-syp/tdT) puncta. Micrographs of 3µm confocal z-stacks show puncta on the medial surface of a soma (top) and a dendrite (bottom). **(D)** Representative reconstruction of excitatory SGN puncta labeled with *Foxg1*^Cre^-dependent syp/tdT (*Foxg1*-syp/tdT; blue) onto a *Thy1* sparsely labeled octopus cell (white). **(E)** Representative reconstruction of inhibitory puncta labeled with *Glyt2*-syp/tdT (green) onto a *Thy1* sparsely labeled octopus cell (white). **(F)** Puncta density for excitatory SGN (*Foxg1*-syp/tdT; black: 10.7 ± 3.0, *n* = 8 cells, 4 mice) and glycinergic puncta (*Glyt2*-syp/tdT; green: 4.2 ± 0.8, *n* = 8 cells, 3 mice) on octopus cell dendrites. Data are presented as mean ± SD. **(G)** Puncta density on somas for excitatory SGN (black: 13.3 ± 2.2, *n* = 8 cells, 4 mice) and glycinergic puncta (green: 1.8 ± 0.1, *n* = 8 cells, 3 mice) and the density along the length of dendrites. Data are presented as mean ± SEM. **(H)** *Top:* Illustration of an octopus cell and the ratio between excitatory SGN puncta and glycinergic puncta. *Inset:* Il-lustration of an octopus cell and the relative innervation densities of excitatory SGNs (blue) and inhibitory puncta (green).

In the mammalian auditory system, precise encoding of broadband timing information begins with the octopus cells of the CNC. Octopus cells are excitatory neurons that bind together co-occurring frequency information on a submillisecond timescale and send this information along one of the parallel ascending pathways in the auditory brainstem. Octopus cells are named for their large-diameter tentacle-like dendrites^15,16^, which are oriented unidirectionally across a tonotopic array of SGNs such that each neuron integrates inputs from a wide range of frequencies^17–20^. SGNs provide the major excitatory inputs onto octopus cells. Biophysically, octopus cells have low input resistances near rest (∼4MΩ)^21,22^, fast time constants (∼200µs)^21–23^, and large low-voltage-activated potassium (∼40nS at rest)^21,23^ and hyperpolarization activated (∼62nS at rest)^21,22^ conductances. Together these properties give octopus cells impressively narrow windows of coincidence detection on the order of 1 millisecond^24–28^. This combination of receiving SGN innervation from broad frequencies and their biophysical specializations establish octopus cells as spectrotemporal coincidence detectors that can reliably encode the timing of complex stimuli, such as the broadband transients found in speech and other natural sounds^21,29,30^. Fittingly, *in vivo* recordings from octopus cells demonstrate their ability to phase lock to broadband transients at rates up to 1kHz^31–33^. Moreover, computational models of octopus cells demonstrate that onset responses are governed by the cell’s biophysical specializations and are, in large part, the result of temporal summation of excitation^34–40^. The simplicity of its connectivity combined with the precision of its temporal computations makes the octopus cell an attractive model for understanding how specialized anatomical and electrophysiological properties contribute to neuronal computations.

Although the octopus cell’s integration of SGN inputs within a narrow time frame enables exceptional coincidence detection, such a model does not explain how other temporal features of sound stimuli are encoded. Indeed, octopus cells encode spectrotemporal sequences within their broadly-tuned response areas, like frequency modulated sounds, that likely require further circuit specializations^41^. Although somatic depolarization can be sufficient to activate an octopus cell^35^, the vast majority of synapses are found on dendrites. Further, SGN inputs are organized tonotopically along octopus cell dendrites, with inputs from high frequency regions located more distal than those from low frequency regions. Dendritic morphology, passive cable properties, active resting membrane properties, and the spatial relationship between synaptic inputs can all impact excitatory post synaptic potential (EPSP) summation as excitation sweeps across the dendritic arbor and towards the soma. This raises the possibility that computations made in the dendrites influence the effective window of coincidence detection by the octopus cell. Such mechanisms could enable flexible processing that is adaptive to the dynamics of the environment without compromising high-fidelity coincidence computations.

Here, we sought to define the circuit mechanisms that allow octopus cells to act as coincidence detectors across time scales. We generated a comprehensive anatomical and physiological map of excitatory and inhibitory synaptic inputs onto octopus cell somas and dendrites and examined how this circuit organization influences octopus cell activation. Through a combination of *in vitro* experiments and computational modeling, we show that somatic summation of excitation is shaped by dendritic inhibition. Thus, octopus cells depend on compartmentalized computations that enable preservation of timing information both at the moment of stimulus onset and within an extended window for evidence accumulation, which is fundamental for the spectrotemporal integration of natural auditory stimuli.

## Results

### The balance of excitatory and inhibitory synapses is different in somatic and dendritic domains

To determine the wiring pattern that drives octopus cell computations, we generated a detailed map of excitatory and inhibitory synaptic inputs (**Fig. 1**). Overall, octopus cells receive abundant excitatory VGLUT1+ innervation from SGNs^42,43^ and sparse inhibitory innervation from glycinergic neurons, as visualized using the glycinergic Cre driver *Glyt2*^Cre^and the Ai34 synaptophysin-tdTomato (syp/tdT) fusion protein reporter (**Fig. 1B**). The *Glyt2-* syp/tdT inhibitory inputs nestle between SGN inputs, especially on octopus cell dendrites (**Fig. 1C**).

Quantification of the number and distribution of presynaptic puncta onto octopus cells revealed marked differences in the ratio of excitation and inhibition in the somatic and dendritic compartments. Since innervation patterns have never been systematically analyzed, we made three-dimensional reconstructions of 16 octopus cells and their excitatory SGN (*n* = 8 cells, 4 mice) and inhibitory (*n* = 8 cells, 3 mice) inputs. Octopus cells were visualized using a *Thy1* reporter and presynaptic puncta were labeled with the syp/tdT reporter driven either by *Foxg1*^Cre^(**Fig. 1D**) or *Glyt2*^Cre^ (**Fig. 1E**). Consistent with qualitative assessment, the density of SGN inputs was higher (10.7 ± 3.0 SGN puncta/100µm^2^) than that of inhibitory inputs (4.2 ± 0.8 puncta/100µm^2^, **Fig. 1F**). Moreover, the relative proportions of excitatory and inhibitory inputs differed in the soma and dendrites (**Fig. 1G**). On somas, SGNs provided dense innervation that continued on the proximal dendrite, then gradually declined with distance from the soma. By contrast, somas received very few inhibitory inputs. On dendrites, inhibitory puncta were evenly distributed. As a result, octopus cells have a strikingly different average ratio of excitatory and inhibitory puncta on the soma (7:1) and on the dendrite (5:2), suggesting that each compartment contributes differentially to the final computation made by the octopus cell (**Fig. 1H**).

### The majority of excitatory synapses on octopus cells are from type Ia SGNs

Although uniformly glutamatergic, SGNs exhibit stereotyped physiological differences in response thresholds that could affect the nature of their inputs onto octopus cells and hence coincidence detection^5,44–46,46,47^. There are three molecularly distinct SGN subtypes, referred to as Ia, Ib, and Ic SGNs, which correlate with previously shown physiological groups^10,48–53^ (**Fig. 2A**). Therefore, we further categorized excitatory inputs based on SGN subtype identity. These can be identified with the presence of *Ntng1*^Cre^-dependent reporter expression^54^ in Ib and Ic SGNs (Ib/c) and its absence in Ia SGNs^50,51,53^, coupled with very low to undetectable levels of calretinin (CR-) in Ic SGNs, and moderate to high levels of calretinin (CR+, CR++) in Ib and Ia SGNs (**Fig. 2B**). As determined using the Ai14 tdTomato (tdT) reporter, *Ntng1*^Cre^-labeled Ib/c SGNs accounted for 60.1 ± 2.6% of the entire population, with 28.5 ± 12.2% Ib SGNs, 31.6% Ic SGNs, and 39.9 ± 2.6% Ia SGNs (**Fig. 2C**: *n* = 1599 neurons, 4 mice; mean ± SD). These proportions matched scRNA-seq estimates (**Fig. 2C**), indicating that this approach provides full coverage. SGN subtype identity was further confirmed by examining the spatial organization of SGN peripheral processes en route to the inner hair cells (IHCs) in the cochlea, with Ia processes positioned deeper than Ib and Ic processes (**Fig 2A, Supp. Fig. 1A-C**).

**Figure 2.**
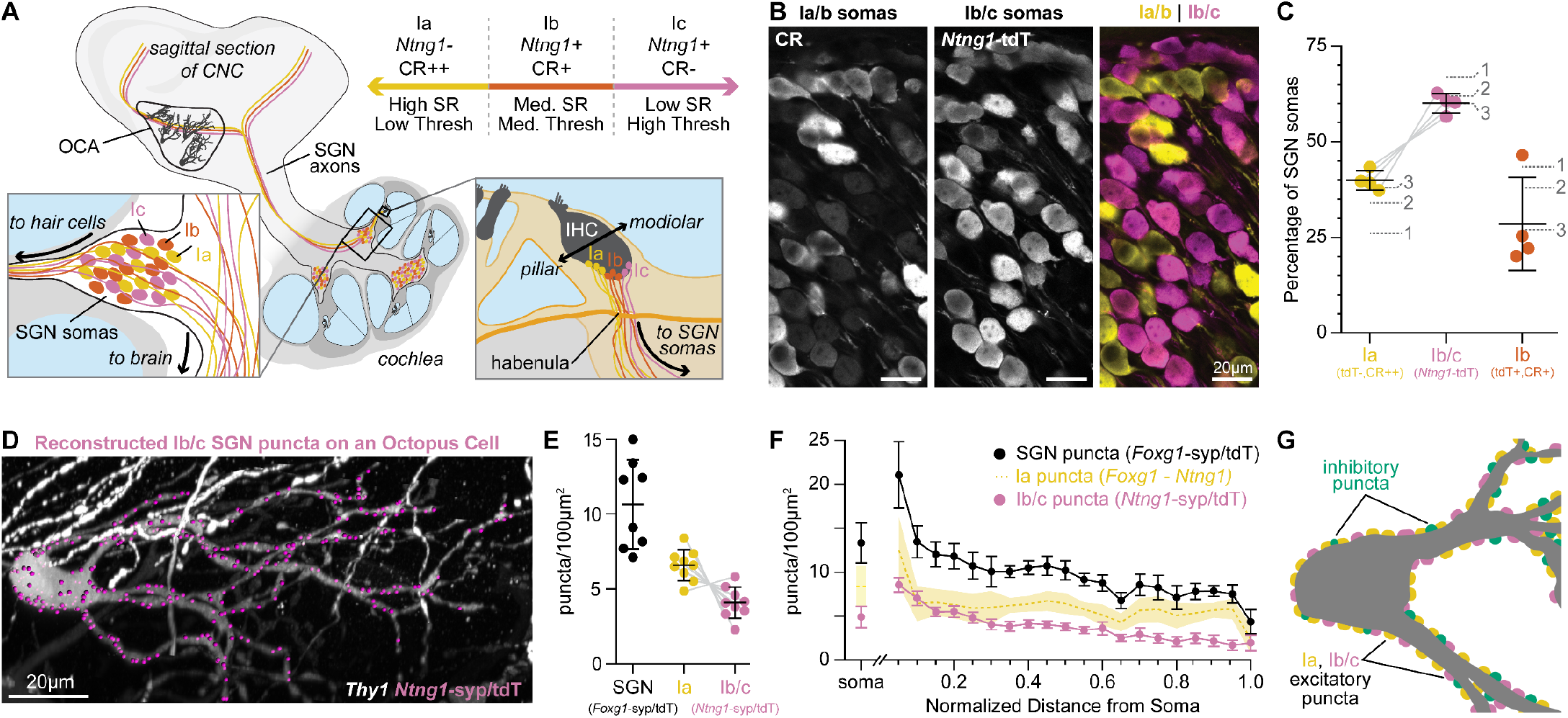
Type Ia SGNs are the primary excitatory contributors to octopus cells. **(A)** Ia (yellow), Ib (orange), and Ic (magenta) spiral ganglion neuron (SGN) axons innervate the cochlear nucleus complex (CNC). SGNs have a continuum of properties organized along the pillar-modiolar axis of inner hair cells (IHCs) and the habenula. *Ntng1-*expressing Ib/c fibers are positioned on the modiolar side (closest to the ganglion). Strongly calretinin immunopositive Ia fibers are on the other side (closest to the pillar cells). This organization correlates with spontaneous rates (SR) and thresholds (thresh.) measured *in vivo*. Somas of all SGN subtypes are found at all tonotopic locations. **(B)** Calretinin (CR) immunolabeling distinguishes SGN subtypes. Ia/b somas label with high (CR++) and medium (CR+) levels of CR, respectively. Ic somas label with low to undetectable levels of CR (CR-). *Ntng1*^Cre^-mediated expression of tdT (*Ntng1*-tdT) labels Ib/c SGNs. **(C)** Ia SGNs (tdT-, CR++) make up 39.9 ± 2.6% of the SGN population. Ib/c SGNs (tdT+) make up 60.1 ± 2.6% of the SGN population. Ib SGNs (tdT+CR+) make up 28.5 ± 12.2% of the SGN population (*n* = 1599 neurons, 4 mice). Data are presented as mean ± SD; individual points represent percent coverage per animal, lines connect measurements from the same animal. Dotted lines indicate percentages from 1: Petitpré et al. 2018, 2: Shrestha et al. 2018, and 3: Sun et al. 2018. **(D)** Representative reconstruction of Ib/c puncta labeled with *Ntng1*^Cre^-dependent syp/tdT (*Ntng1*-syp/tdT; magenta) onto a *Thy1* sparsely labeled octopus cell (white). **(E)** Puncta density for all SGNs (*Foxg1*-syp/tdT; black: data from Fig. 1F), Ia SGNs (*Foxg1 - Ntng1*; yellow: 6.6 ± 1.0), and Ib/c SGNs (*Ntng1*-syp/tdT; magenta: 4.1 ± 1.0, *n* = 9 cells, 5 mice) along the total dendritic length. Ia density was calculated by subtracting Ib/c density from total SGN density; lines connect measurements from the same reconstruction. Data are presented as mean ± SD. **(F)** Puncta density on somas for all SGNs (*Foxg1*-syp/tdT; black: data from Fig. 1E), Ia SGNs (*Foxg1 - Ntng1*; yellow: 8.4 ± 2.3), and Ib/c SNGs (*Ntng1*-syp/tdT; magenta: 4.9 ± 1.2, *n* = 9 cells, 5 mice) and the density along the length of dendrites. Data are presented as mean ± SEM. **(G)** Illustration of an octopus cell and the relative innervation densities of Ia SGNs (yellow), Ib/c SGNs (magenta), and inhibitory puncta (green).

Within the ventral cochlear nucleus (VCN), where octopus cells reside, *Ntng1*^Cre^ labeling was restricted to SGN central axons (**Supp. Fig. 1D-E**). There is sparse labeling in the deep layer of the dorsal cochlear nucleus (DCN) and strong labelling throughout the thalamus, hippocampus, and cortex (**Supp. Fig. 1D-E**). It is unlikely that the *Ntng1*^Cre^ labeled cells outside of the cochlea make synapses with octopus cells. When *Ntng1*^Cre^ driven tdT is co-expressed with *Foxg1*^Flp^ driven EYFP, all tdT-labeled axons in the VCN also expressed EYFP, suggesting that all our *Ntng1*^Cre^ labeled inputs are a part of the *Foxg1*^Flp^ labeled population, which are very likely to only be from SGNs in the cochlea. Additionally, in the octopus cell area, CR+ SGN central axons were segregated from *Ntng1*-tdT labeled central axons, consistent with the moderate to undetectable levels of CR in *Ntng1*-tdT SGN somas in the periphery (**Supp. Fig. 2F**). Thus, *Ntng1*^Cre^-driven expression of syp/tdT is an appropriate tool for mapping subtype-specific connectivity onto octopus cells. Reconstruction of *Ntng1*^Cre^-labeled Ib/c puncta (**Fig. 2D**) demonstrated that octopus cells are dominated by inputs from Ia SGN fibers, which have the lowest response thresholds and highest rates of spontaneous activity. Octopus cell dendrites received 4.1 ± 1.0 puncta/100µm^2^ from Ib/c SGNs (**Fig. 2E**, magenta: *n* = 9 cells, 5 mice; mean ± SD), accounting for 38% of the total SGN density. Given that *Ntng1*-tdT+ cells account for 60.1% of the SGN population (**Fig. 2C**, magenta), Ib/c inputs were underrepresented on octopus cells. Octopus cells receive similarly low innervation from Ic inputs (**Supp. Fig. 2G-I**: *n* = 6 cells, 2 mice), as estimated from the degree of sparse labeling of Ic axons achieved by *Myo15*^iCre^-driven reporter expression (**Supp. Fig. 2A-F**) and the expected proportion of Ic SGNs in the ganglion (**Supp. Fig. 2F**). By contrast, Ia SGNs, which comprise ∼40% of the total population (**Fig. 2C**, yellow), accounted for 62 ± 9.7% of SGN synapses on octopus cells (**Fig. 2E**, yellow: 6.6 ± 1.0 puncta/100µm^2^). All three subtypes showed the same overall distribution from the soma to the distal dendrite (**Fig. 2F**). Together, excitatory and inhibitory puncta densities in the innervation maps indicate the average octopus cell receives ∼1035 SGN synapses (642 Ia SGN, 393 Ib/c SGN) and ∼354 inhibitory synapses. Additionally, 83% of the synapses were on the dendrites, suggesting a critical role in the octopus cell computation. Octopus cell reconstructions showed the same basic wiring patterns regardless of where each cell was positioned in the octopus cell area. The tonotopic position of all reconstructed octopus cell somas was estimated in 3D reconstructions aligned to a normalized CNC model of tonotopy^55^. Octopus cells had similar morphologies (**Supp. Fig. 3E-G**) and patterns of synaptic innervation (**Supp. Fig. 3H-M**) regardless of where they were positioned along the tonotopic axis. Thus, Ia SGN synapses are the primary inputs to both the soma and dendrites of octopus cells. Additionally, the whole-neuron wiring diagram identifies a dendritic domain where inhibitory synapses are approximately equal in number to the Ib/c excitatory synapses from the periphery.

### SGN inputs to octopus cells facilitate at high stimulation frequencies

Whether or not an octopus cell responds to its inputs depends on when and how EPSPs travel to and then summate in the soma. To determine if SGN subtypes transmit information differently to their central targets, we performed *in vitro* whole-cell current clamp recordings of octopus cells (**Fig. 3A**) while using Cre-dependent Channelrhodopsin-2 (ChR2) to evoke EPSPs from all SGNs (*Foxg1*-ChR2) or only Ib/c SGNs (*Ntng1*-ChR2). Octopus cells have short action potentials (∼5-15mV) that resemble their large, well-timed EPSPs^25,56^. Therefore, we evoked small EPSPs that were below spike threshold and distinguishable from action potentials using a phase plot analysis. Trains of ChR2-evoked EPSPs, ranging from 5 to 50Hz, in both the total SGN population (**Fig. 3B**, black: *n* = 8 cells, 5 mice) and the Ib/c SGN population (**Fig. 3B**, magenta: *n* = 7 cells, 6 mice) exhibited no differences in paired-pulse plasticity at any frequency of stimulation (p > 0.35 at all interstimulus intervals, Tukey’s HSD), although the Ib/c population exhibited higher variability than the total SGN population (at 20ms: SD = 0.11, SD = 0.24, respectively).

**Figure 3.**
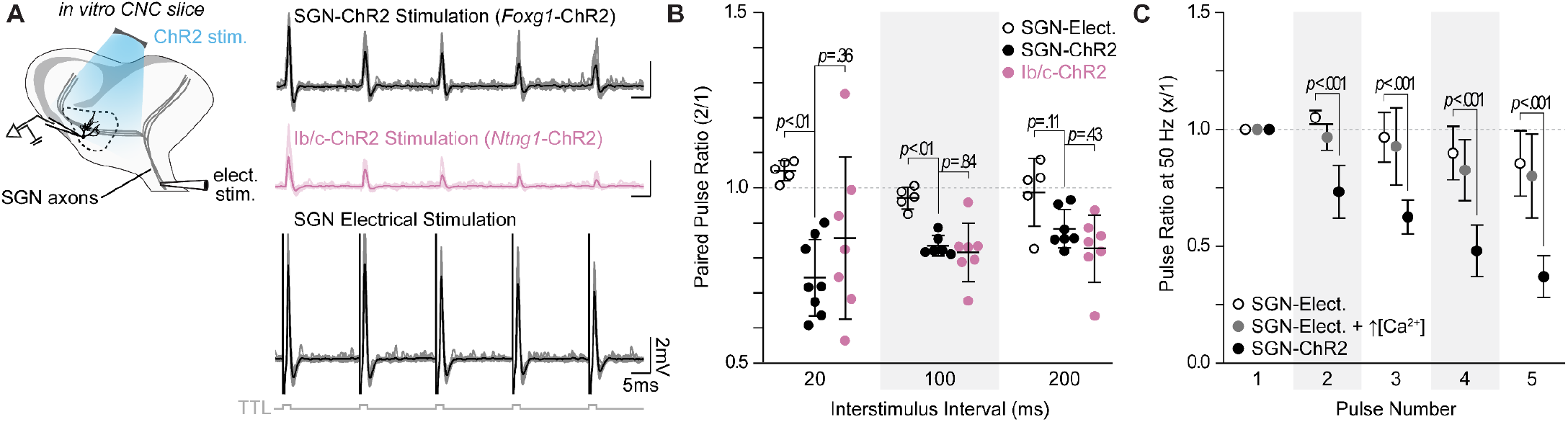
SGN subtype inputs to octopus cells do not differ in short term plasticity. **(A)** Illustration of the experimental paradigm and representative EPSP traces recorded during *in vitro* whole-cell current-clamp recordings of octopus cells. Spiral ganglion neuron (SGN) central axon stimulation methods included electrical stimulation or full-field, light-evoked activation of *Foxg1-* ChR2 or *Ntng1*-ChR2 SGNs. TTL trigger pulses are shown in gray. **(B)** Paired pulse ratios for electrically stimulated SGNs (open circles: *n* = 5 cells, 3 mice), ChR2 stimulated SGNs (*Foxg1*-ChR2; black: *n* = 8 cells, 5 mice), and ChR2 stimulated Ib/c SGNs (*Ntng1*-ChR2; magenta: *n* = 7 cells, 6 mice) at three interstimulus intervals. With electrical stimulation, SGN inputs to octopus cells were stable and exhibited slight facilitation at 50 Hz (20ms interstimulus interval). ChR2 stimulation caused paired pulse depression not seen with electrical stimulation. Data are presented as mean ± SD. Each data point represents the average paired pulse ratio for a cell. *p* values from ANOVA and subsequent Tukey HSD test are reported for comparisons between methods of SGN activation (electrical and ChR2) and SGN subpopulation composition within method of activation (SGN-ChR2 and Ib/c-ChR2). Welch’s ANOVA was used for comparisons at 20ms interstimulus interval (50Hz) as data in this condition did not meet the homogeneity of variance assumption. **(C)** Pulse ratios at 50Hz for electrically stimulated SGNs with physiological 1.4mM Ca^2+^ ACSF (open circles: *n* = 5 cells, 3 mice), electrically stimulated SGNs with 2.4mM Ca^2+^ ACSF (grey: *n* = 3 cells, 2 mice) and ChR2 stimulated SGNs with physiological ACSF (*Foxg1*-ChR2; black: *n* = 8 cells, 5 mice). Data are presented as mean ± SD. *p* < 0.001 from ANOVA and subsequent Tukey HSD test for all comparisons between methods of SGN activation (electrical and ChR2). There were no statistically significant differences for all comparisons under 1.4mM and 2.4mM Ca^2+^ (*p* > 0.100, ANOVA).

ChR2-evoked synaptic responses are known to undergo synaptic depression^57,58^. To determine if the paired-pulse depression measured in ChR2-stimulated experiments was physiological, we used electrical stimulation to evoke EPSPs from SGNs (**Fig. 3B**). Electrically-evoked EPSPs had higher paired-pulse ratios than ChR2-evoked EPSPs and were mildly facilitating at short (20ms) intervals (**Fig. 3B-C**, open circles: *n* = 5 cells, 3 mice), consistent with an octopus cell’s ability to respond reliably to click trains *in vivo*^17,19,32,59^. In contrast, previous results using electrical stimulation demonstrated short-term depression of SGN inputs to octopus cells^60,61^. However, these experiments were carried out in the presence of higher, non-physiological levels of extracellular calcium. We repeated paired-pulse plasticity experiments with non-physiological calcium concentrations (2.4mM) and similarly found that electrically-evoked EPSPs from SGNs resulted in short-term depression at 50Hz of electrical stimulation (**Fig. 3C**, grey: *n* = 3 cells, 2 mice), though not to the degree observed when using full-field, ChR2-evoked inputs.

### Glycine evokes inhibitory post synaptic potentials that are occluded by a low input resistance

Given the high density of inhibitory synapses on octopus cell dendrites, we considered the possibility that somatic and dendritic compartments contribute differently to the temporal computation made by octopus cells. A role for inhibition has never been incorporated into octopus cell models as previous efforts failed to reveal physiological evidence of functional inhibitory synapses either *in vitro*^25,56,62^ or *in vivo*^41^. Similarly, we did not observe light-evoked (*Glyt2*-ChR2) inhibitory post synaptic potentials (IPSPs) in octopus cell somas during whole-cell current clamp recordings from P30-45 mice. Since inhibitory synapses are located primarily on octopus cell dendrites, we posited that their voltage spread to the soma is limited given the extremely low input resistance of octopus cells.

To decrease electrotonic isolation of the dendrites and increase input resistance, we pharmacologically reduced conductances from voltage-gated potassium (Kv) and hyperpolarization-activated cyclic nucleotide-gated (HCN) channels using 100µM 4-Aminopyridine (4-AP) and 50µM ZD 7288 (ZD). This cocktail increased octopus cell membrane resistance (**Fig. 4A**) and hyperpolarized the resting membrane potential ∼8-10mV. To compensate for this change, membrane potentials were adjusted to within 3mV of the original resting membrane potential with a holding current. To isolate inhibition, AMPA receptor activation was blocked using 15µM 2,3-dioxo6-nitro-7-sulfamoyl-benzo[f]quinoxaline (NBQX). Consistent with our hypothesis, the increase in input resistance unveiled light-evoked IPSPs in recordings from octopus cell somas (**Fig. 4B**). Additionally, these IPSPs were fully abolished by bath application of 500nM strychnine (STN) (**Fig. 4C**), confirming the presence of functional glycinergic inhibitory synaptic transmission onto octopus cells.

**Figure 4.**
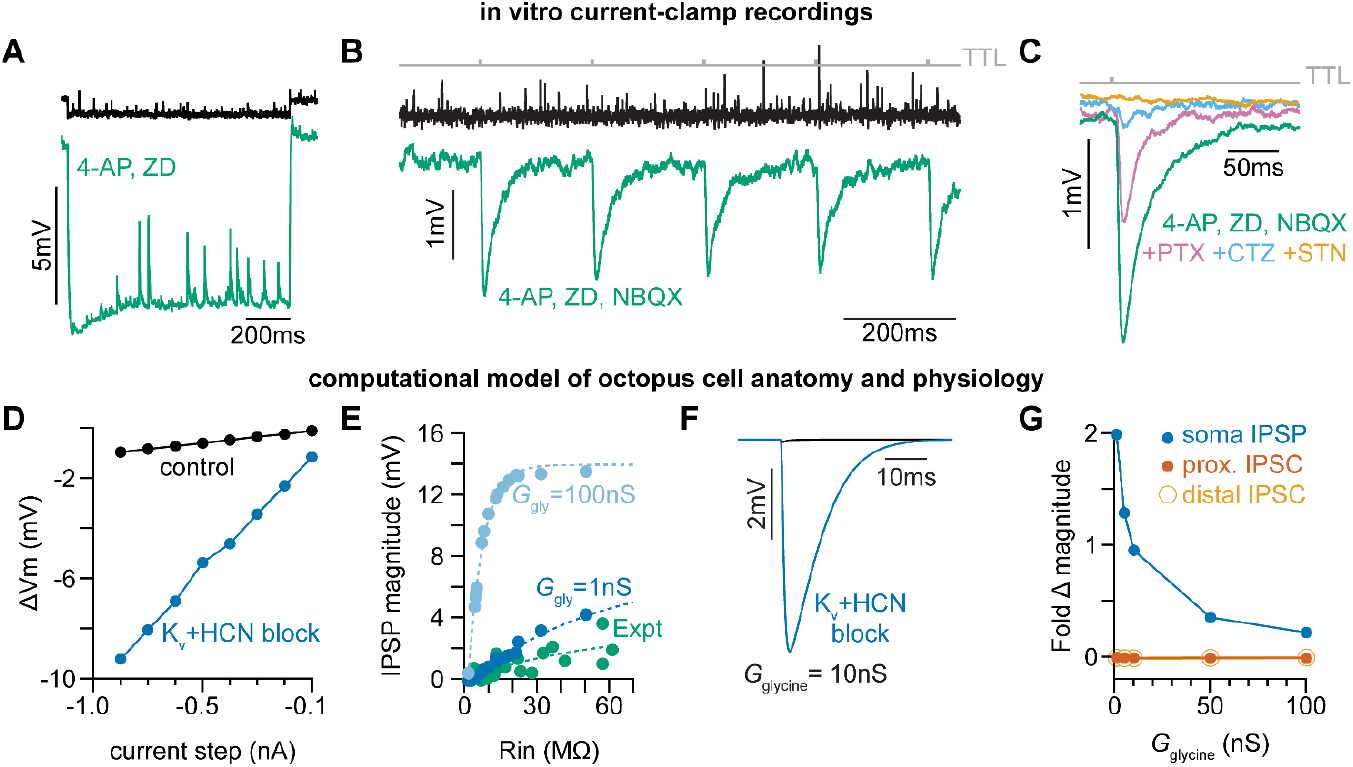
Octopus cells receive glycinergic inhibitory post synaptic potentials. **(A)** Voltage responses to a -200pA current injection. This representative neuron hyperpolarized 0.7mV (black) in control conditions. After bath application of 100µM 4-Aminopyridine (4-AP) and 50µM ZD 7288 (ZD), hyperpolarizing responses to the same -200pA current injection increased to 8.8mV at steady state (green). **(B)** Postsynaptic responses to ChR2 stimulation of glycinergic terminals (*Glyt2*-ChR2) with a 5Hz train (gray) of 1ms full-field blue light pulses before (black) and after bath application of 100µM 4-AP, 50µM ZD, and 15µM NBQX (green: *n* = 9 cells, 8 mice). Increased input resistance reveals inhibitory potentials that are difficult to detect. **(C)** Postsynaptic responses to *Glyt2*-ChR2 stimulation after bath application of 100µM 4-AP, 50µM ZD, and 15µM NBQX (green), with further sequential addition of 20µM picrotoxin (PTX, pink), 100µM cyclothiazide (CTZ, blue), and 500nM strychnine (STN, orange; *n* = 6 cells, 5 mice). **(D)** Change in membrane voltage in response to hyperpolarizing somatic current steps in a morphologically and biophysically realistic model of octopus cells before (black) and after removal of voltage-gated potassium (Kv) and hyperpolarization-activated cyclic nucleotide-gated (HCN) channels (blue). As in *in vitro* current-clamp recordings *(A)*, removing Kv and HCN channels increased the magnitude of voltage responses (ΔVm) to hyperpolarizing current. **(E)** IPSP magnitude in experimental data (green) and the model (dark and light blue) as a function of input resistance. In somatic measurements, IPSP size increases with input resistance. Modeled IPSPs are shown for two conductance levels (1nS, dark blue; 100nS light blue). **(F)** IPSPs measured at the soma of a modeled octopus cell before (black) and after removal of Kv and HCN channels (blue). As in *in vitro* current-clamp recordings, this allows for somatic IPSP detection. **(G)** Fold change in magnitude of soma-measured IPSPs (blue) or dendrite-measured IPSCs at proximal (dark orange) and distal (light orange, open circles) dendritic locations after removal of Kv and HCN channels. Kv and HCN block increases the magnitude of soma-measured IPSPs. The size of dendritic IPSCs are not changed with Kv and HCN block.

To determine the types of glycinergic receptors contributing to IPSPs, we pharmacologically blocked subsets of glycine receptors (**Fig. 4C**). IPSPs were reduced upon addition of 20µM picrotoxin (PTX), which blocks homomeric glycine receptors^63–65^. Sequential addition of 100µM cyclothiazide (CTZ), which blocks α2-containing homomeric and heteromeric glycine receptors^66,67^, nearly abolished the remaining IPSPs, and responses were fully abolished with further application of 500nM STN. These results indicate that relevant glycine receptors include both large conductance extrasynaptic β-subunit lacking homomeric receptors and synaptically localized α2β receptors with slower kinetics^68–70^.

To confirm whether the electrical confinement of IPSPs to the dendrites is consistent with our understanding of octopus cell biophysics, we developed an improved biophysically and anatomically accurate model of octopus cells based on our findings (**Supp. Fig. 4**)^30,71^. Passive and active properties of the model were first aligned with experimental data using a scaling factor (scl), which adjusts the maximum conductance of voltage-gated potassium (Kv) 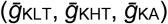 and hyperpolarization-activated cyclic nucleotide-gated (HCN) 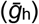 channels (**Supp. Fig. 4A-B**). To further align the model with experimental data, we adjusted input resistance (*R*_*N*_) and specific membrane resistance (*R*_*m*_) to match experimentally measured input resistance under physiological conditions and with Kv and HCN block (**Supp. Fig. 4D-E**). We further examined the influence of leak, Kv, and HCN conductances on the neuron’s input resistance. In a passive scenario, where leak, Kv, and HCN conductances are set to zero (*g*_leak_*=0*; *g*_Kv_=0; *g*_h_=0), the input resistance scaled proportionally with specific membrane resistance (**Supp. Fig F**, grey). However, with active Kv and leak conductances, *R*_*N*_ was constrained and showed only a slight increase relative to *R*_*m*_, suggesting attenuation of PSPs during their propagation to the soma (**Supp. Fig. 4F-G**). As in our current-clamp recordings (**Fig. 4A**), removal of Kv and HCN conductances in the model changed the input resistance and current-voltage relationship of the neuron, resulting in reduced electrotonic isolation (**Supp. Fig. 4C, Fig. 4D**, blue). We next tuned the glycine conductance (*G*_gly_) to match experimental results (**Fig. 4E)**. Activation of large (10nS) dendritic glycinergic conductances induced negligible hyperpolarizing voltage changes in the model (**Fig. 4F**, black). With increased input resistance and reduced electrotonic isolation, dendritic IPSPs measured at the soma were detectable, consistent with our *in vitro* recordings (**Fig. 4F**).

While blocking Kv and HCN allowed us to reveal IPSPs at the soma, it is possible that reduced electrotonic isolation does not entirely explain the increase in somatically-measured IPSP amplitude. Changes in driving force could increase the magnitude of synaptic currents and therefore increase the magnitude of experimentally measured synaptic potentials. We used the model to explore how changes in membrane potential under Kv and HCN block could impact the magnitude of post synaptic potentials. We simulated dendritic glycinergic conductances over a range of values and in the presence of blocked Kv and HCN channels (**Supp. Fig. 4H**). Kv and HCN block and the resulting change in input resistance increased the magnitude of soma-measured IPSPs for all glycine conductances (**Fig. 4G**, blue). The size of inhibitory currents measured in the dendrites did not change as a result of the hyperpolarization induced by Kv and HCN block regardless of synaptic location on the dendrites or the magnitude of the glycine conductance (**Fig 4G**, dark orange, light orange; **Supp. Fig 4I-J**). These observations suggest that the IPSPs are difficult to detect in physiological conditions because of severe attenuation during transmission along dendrites to the soma. This is further supported by marked changes in transfer impedance (ZT), a measure of signal transmission efficiency, measured with and without Kv and HCN block (**Supp. Fig. 4K**). Together, results from the model provide evidence of functional glycinergic synaptic transmission that is difficult to detect with *in vitro* somatic recordings.

### Inhibition decreases the magnitude and advances the timing of dendritic SGN inputs

SGN synapses onto octopus cell dendrites are arranged tonotopically, with higher frequency SGNs from the base of the cochlea terminating on distal dendrites and lower frequency SGNs from more apical positions terminating proximally (**Fig. 1A**). This organization has been proposed to re-synchronize coincidentally firing SGNs that are activated at slightly different times due to the time it takes for the sound stimulus to travel from the base to the apex of the cochlea^30^. To test how inhibition in the dendrites shapes coincidence detection, we first used our model to explore the influence of simultaneous activation of inhibitory and excitatory synapses at varying locations along the dendritic tree^72,73^. We modelled how somatically recorded EPSPs are affected by the location of inhibition by moving the site of excitation relative to inhibitory synapses placed either on proximal or distal dendrites (**Fig. 5A, Supp. Fig. 5**). In our model, inhibitory synapses that are located proximally to excitation (**Supp. Fig. 5A**) had less of an influence on excitation recorded at the soma compared to those inhibitory synapses located distally to excitation (**Supp. Fig. 5B**). We analyzed the effect of inhibitory synapse location on somatically measured EPSPs using varying excitatory and inhibitory synaptic weight values (**Supp. Fig. 5C-F**) and determined that inhibitory conductances (*G*_Gly_) between 6 and 10nS produced values within experimental ranges. For both inhibition proximal to excitation (**Fig. 5B-C**) and inhibition distal to excitation (**Fig. 5D-E**), the model predicted that inhibition reduces EPSP amplitude and accelerates EPSP peak timing at the soma. Thus, the presence of inhibition could modulate EPSP timing in dendritic compartments during continuous auditory stimuli, when inhibition can be recruited after the onset of a sound and thus allow for adaptable temporal processing during the sound’s duration.

**Figure 5.**
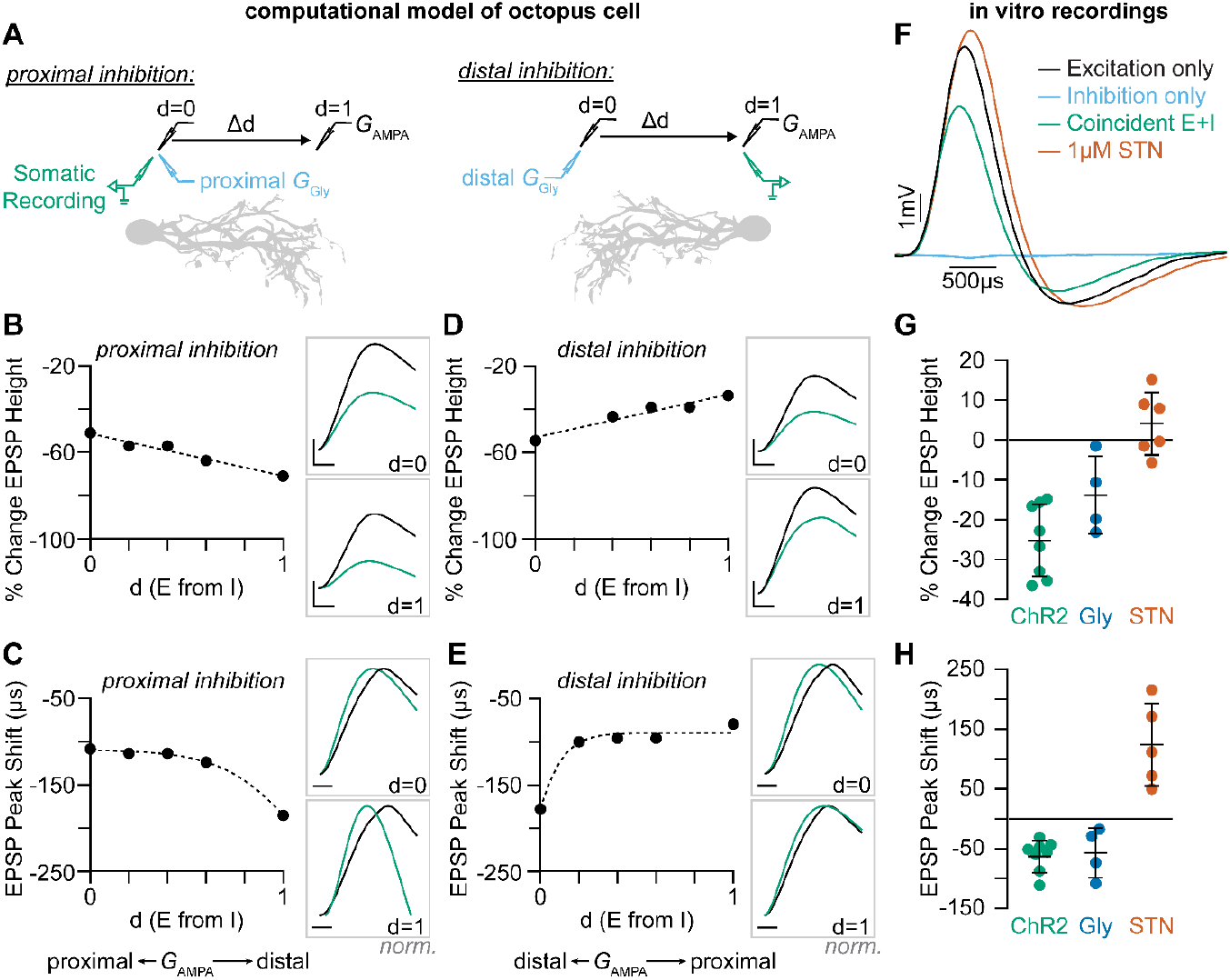
Coincident excitation and inhibition on octopus cell dendrites advances EPSP peak times. **(A)** The impact of distance between excitatory and inhibitory synapses was measured in a computational model of octopus cells. Inhibitory synapses were placed either proximally or distally to excitation. Excitatory synapses were placed at varying locations along the dendritic arbor to change the anatomical distance (Δd) where d=0 is the location of inhibition (*G*_Gly_) and d=1 is the condition where excitation (*G*_AMPA_) and inhibition are maximally separated. EPSPs were measured at the soma in all conditions (green). **(B-E)** Quantification of the percent change in soma-measured EPSP magnitude *(B, D; G*_AMPA_=5nS) and the shift in EPSP peak timing *(C, E; G*_AMPA_=2nS) in models of proximal *(B-C)* and distal *(D-E)* inhibition. Inhibitory conductances (*G*_Gly_) between 6 and 10nS produced values within experimental ranges. Example traces show EPSPs with (green) and without (black) inhibition at d=0 and d=1. Distal dendrites (d=1) have higher local input resistance and lower IPSP attenuation due to the sealed end. Inset scale bars are 1mV, 200ms. **(F-H)** Coincident stimulation of excitation and inhibition changes EPSP shape. (*F)* Representative responses to independent stimulation of excitatory spiral ganglion neurons (SGNs; black), independent stimulation of inhibitory inputs (light blue), coincident stimulation of both excitation and inhibition (green), and independent stimulation of excitatory SGNs with the addition of 1µM strychnine (STN). Quantification of the percent change in EPSP height *(G)* and the shift in EPSP peak timing *(H)* during coincident *Glyt2*-ChR2 activation of inhibitory inputs (green: *n* = 8 cells, 6 mice), bath application of 25µM glycine (dark blue: *n* = 4 cells, 3 mice), and bath application of 1µM STN (orange: *n* = 5 cells, 4 mice). Activation of glycinergic receptors during excitation decreased EPSP heights and advanced EPSP peaks. Blocking of tonically active glycine receptors slowed and delayed EPSPs. Data are presented as mean ± SD. Markers represent the average quantification for a cell.

To directly test if the prediction that temporally coincident excitation and inhibition affects the timing and amplitude of excitatory SGN inputs as they travel towards the octopus cell soma, we coincidently activated SGNs and glycinergic inputs *in vitro*. In these experiments, the octopus cell properties were not altered pharmacologically and inhibition was undetectable or only visible with averaging over many sweeps (**Fig. 5F**, blue). When synaptic inhibition was evoked together with excitation (**Fig. 5F**, green), EPSPs recorded in the soma were smaller and faster than when excitation was evoked alone (**Fig. 5F**, black: *n* = 8 cells, 6 mice). ChR2-evoked inhibition decreased EPSP heights by 25.2 ± 9.0% (**Fig. 5G**, green) and shifted the peak of EPSPs forward 57.5 ± 26µs (**Fig. 5H**, green). This effect was mimicked by bath application of 25µM glycine (**Fig. 5G-H**, blue: *n* = 4 cells, 3 mice). Further, bath application of 1µM STN had the opposite effect, resulting in larger EPSPs and delayed peak times (**Fig. 5F-H**, orange: *n* = 5 cells, 4 mice). These findings suggest that the timing of EPSP arrival in the soma may be shaped both by tonically active glycine channels and the release of synaptic glycine onto the dendrite. Of note, many SGNs also terminate on the octopus cell soma, where inhibition is minimal. This suggests that the octopus cell’s ability to act as a coincidence detector depends on two stages of compartmentalized computations, one in the dendrite that combines excitation and inhibition to provide important information about which frequencies co-occur in a complex sound stimulus and one in the soma that is restricted by the rigid temporal summation window for coincidence detection. Together with the electrotonic properties of the octopus cell and the dominance of low threshold, low jitter Ia SGN inputs, these combined computations can enable reliable coincidence detection and cross-frequency binding needed for perception of sound.

## Discussion

Coincidence detection plays a critical role in many cognitive and perceptual processes, from the ability to localize sound to the binding of auditory and visual features of a common stimulus. Depending on the computation, the temporal window for integration can range widely, thereby requiring circuitry with distinct anatomical and physiological properties. Here, we describe a two-domain mechanism for coincidence detection that can detect co-occurring frequencies with different degrees of precision. By mapping and selectively activating synaptic inputs onto octopus cells both *in vitro* and in a computational model, we revealed that compartmentalized dendritic nonlinearities impact the temporal integration window under which somatic coincidence detection computations are made. The arrival of many small, reliable excitatory inputs (**Fig. 3**) from low-threshold SGNs (**Fig. 2**) is continuous throughout an ongoing stimulus. We demonstrate that glycinergic inhibition to octopus cell dendrites (**Fig. 1**) can shift the magnitude and timing of SGN EPSPs as they summate in the soma (**Fig. 4**,**5**). The narrow window for coincidence detection computations allows the octopus cell to respond with temporal precision using momentary evidence provided by SGNs at the onset of the stimulus. We propose that, as a stimulus persists, inhibition onto octopus cell dendrites can adjust EPSPs before they arrive at the soma for the final input-output computation. This allows the cell to make an additional computation with a slightly longer window for evidence accumulation without compromising the accuracy of the somatic computation (**Fig. 6**).

**Figure 6.**
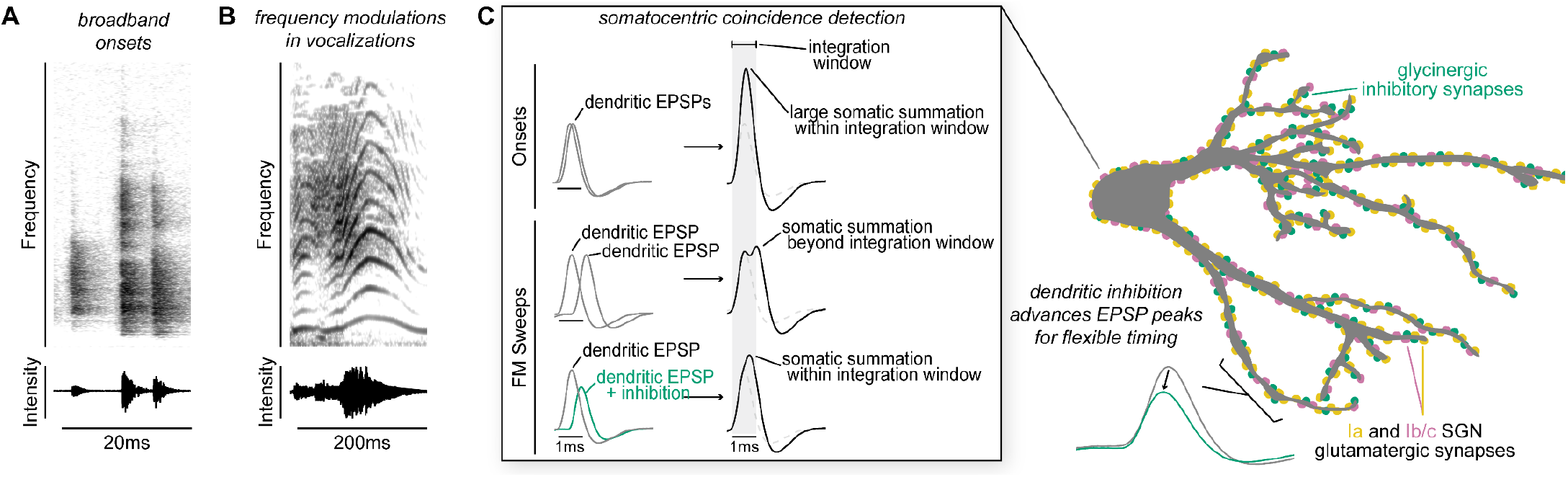
Proposed model of flexible dendritic timing for precise somatic coincidence detection. **(A-B)** Spectrograms show how co-occurring frequencies in broadband onsets (shown for a clicking sound, *A*) and frequency modulations (shown for human squeaking, *B*), change in their strength (top) and total intensity (bottom) on different time scales. Despite the timing differences, both kinds of stimuli are hypothesized to result in somatic summation during the octopus cell’s spike integration window. **(C)** Excitation must summate at the soma of octopus cells within a narrow time window (∼1ms) to achieve a depolarization rate rapid enough to trigger action potentials. Summation of excitation alone accounts for responses to sound onsets (inset, top). During stimuli that require summation over a longer time period, such as frequency modulated sweeps, synaptic inhibition can accelerate EPSPs as they travel along octopus cell dendrites towards the soma for coincidence detection computations (inset, bottom). We propose a mechanism for preferential processing of a subset of excitatory inputs where selective temporal advancement of a subset of EPSPs by local inhibition could expand the effective window for coincidence detection at the soma.

As coincidence detectors in the auditory system, octopus cells are faced with the challenge of recognizing complex sounds that include many frequencies that co-occur from the beginning to the end of the stimulus. As shown *Flexible Coincidence Detection in the Auditory System* by *in vivo* recordings^18,41^, octopus cells respond well to cues that include complex spectrotemporal patterns, including frequency modulations beyond the onset of the stimulus^41^. Given that the auditory environment is filled with overlapping sound stimuli, such responses presumably allow the octopus cell to encode which frequencies belong to which sound. Our data thus support the role of octopus cells beyond simple onset coincidence detectors that rely solely on the temporal summation of excitation. The results suggest that, in addition to high Kv and HCN conductances at rest, the addition of dendritic inhibition transforms the magnitude and timing of SGN signals as they arrive in the cell body, which may expand the response selectivity of an octopus cell and allow them to become a slightly leakier integrator that can accumulate evidence beyond onsets. Although, this inhibition is difficult to detect because of shunting, our data demonstrate that it is both present and impactful.

As well as needing to work beyond onsets, an effective coincidence detector in the auditory system must also function reliably across a range of sound intensities. Intensity information is encoded by the number and types of SGNs that are activated in the cochlea. The Ia, Ib, and Ic molecular subtypes defined in mouse^50,51,53^ broadly correspond to the anatomically and physiologically defined subtypes described across species^52,74^. We find that the majority of inputs onto octopus cells come from Ia SGNs, which most closely correspond to the low-threshold, high-spontaneous rate population. Consistent with this result, single-unit SGN recordings in cats demonstrated a bias towards low-threshold, high-spontaneous rate axon collaterals in the octopus cell area^44^. Low-threshold SGNs are also characterized by short first spike latencies and low temporal jitter^75–78^. A hallmark of the octopus cell is the fact that it only fires action potentials when many SGN inputs are activated within a narrow period of time^79^. The presence of many low-threshold and temporally precise inputs on the octopus cell may help ensure that coincidence detection still works reliably for quiet sounds. Further, Ia inputs onto octopus cells do not exhibit paired-pulse depression, similar to low levels of depression seen in Ia inputs to bushy cells^80^. The presence of SGN inputs without paired-pulse depression could be beneficial for encoding sustained auditory signals. Finally, while Ia SGNs are over-represented, Ib and Ic inputs are also present. Since precise, low-threshold SGN responses can be saturated by background noise such that responses to relevant stimuli are masked^10,75,78,81^, recruitment of higher threshold SGNs at higher sound intensities may compensate for this tradeoff.

The presence of inhibitory inputs onto dendrites is a fundamental feature of the nervous system and, in other systems, contributes to a neuron’s final computation. For example, direction selectivity computations in dendrites of retinal cells require excitation-inhibition interactions in dendritic compartments^82,83^. In pyramidal cells of the cortex and hippocampus, the spatial distribution of inhibition impacts dendritic non-linearities in a branch-selective manner^73,84–88^. However, octopus cells do not share all mechanisms for dendritic computation seen in cortical and hippocampal neurons. Resting conductances and low-threshold potassium conductances may suppress voltage-gated calcium spikes and attenuate the magnitude of action potentials as they backpropagate through the soma and dendritic tree^21,89–91^. Therefore, subthreshold dendritic integration of coincident excitation and local inhibition may be the primary computation that occurs in octopus cell dendrites before action potential generation near the soma.

Although this work uncovers a role for inhibition, a deeper understanding of octopus cell computations will require determining what information is carried by inhibitory inputs. Despite the well-established presence of presynaptic glycinergic puncta in the octopus cell area^92–95^ and glycinergic receptor expression in octopus cells^96–100^, it is unknown where glycinergic innervation originates. Local neurons within the CNC could provide inhibition; however it remains unclear whether octopus cells receive connections from D-Stellate^56^, L-Stellate^101^, or tuberculoventral cells^102^. Outside of the CNC, terminal degeneration experiments in cats suggested the superior periolivary nucleus (SPON) and the ventral division of the lateral lemniscus (VNLL) as potential sources of descending inhibition to the octopus cell area^103^. Octopus cells provide excitatory input to both the SPON^104–108^ and the VNLL^32,109–113^, raising the possibility of feedback inhibition from the auditory brainstem as a circuit mechanism that elongates temporal summation windows during ongoing stimuli. Such descending feedback inhibition is not rapid enough to prevent or alter the characteristic octopus cell onset response, but could change the effective coincidence detection window as the stimulus continues or limit the duration of the response. Future studies will be required to identify the source of inhibition and its organization within dendritic compartments. If inhibitory inputs tonotopically match the local, narrowly-tuned dendritic SGN inputs, it is possible that frequency-matched inhibition could influence spectral selectivity or feature extraction. On the other hand, broadly tuned inhibition could reduce depolarization block or serve as a temporal milestone that signals gaps or offsets. Further characterization of *in vivo* octopus cell responses in complex sound environments may clarify the effect of noise on signal detection and reveal additional features of this cell’s contributions to perception of the auditory world.

## Acknowledgements

We thank Nace Golding (University of Texas at Austin), Matthew McGinley (Baylor College of Medicine), Philip Joris (KU Leuven), and Bernardo Sabatini (Harvard Medical School) for helpful discussions and feedback. Sadie Quinn, Lucy Lee, and Ryan Merrow provided valuable technical assistance. Bruce Bean (Harvard Medical School) generously provided access to electrophysiological equipment. *Slc6a5*^tm1.1(Cre)Ksak^, *Ntng1*^em1(Cre)Kfra^, and *Myo15a*^tm1.1(Cre)Ugds^mice were kindly provided by Wade Regehr (Harvard Medical School), Fan Wang (Massachusetts Institute of Technology), and Stefan Heller (Stanford). We thank Rigoberto Ramirez, Tenzin Paljorwa, and Edgar Ramirez for animal care support. We are grateful to the Neurobiology Imaging Facility (NIF) for software availability and to the HMS Research Instrumentation Core for the design and fabrication of temperature regulation equipment. This work was supported by grants from the BRAIN Initiative 1R01NS118402 to L.V.G., the National Institute on Deafness and Other Communication Disorders 5R01DC009223 to L.V.G. and 1F32DC020070 to L.J.K, the William Randolph Hearst Fund to L.J.K, and the Broad Institute’s Stanley Center for Psychiatric Research to S.H. and G.F.

## Author Contributions

Conceptualization, L.J.K. and L.V.G.; Methodology, S.H.; Formal Analysis, L.J.K and S.H.; Investigation, L.J.K, S.M., and S.S.; Resources, G.F.; Writing, L.J.K. and L.V.G.; Visualization, L.J.K. and S.H.; Supervision, G.F. and L.V.G.; Funding Acquisition, L.J.K, G.F., and L.V.G. Declaration of Interests

## Materials and Methods

### Animal Use and Transgenic Mouse Lines

All procedures were approved by and conducted in accordance with Harvard Medical School Institutional Animal Care and Use Committee. Male and female mice (*Mus musculus*) were bred on a C57BL/6 background at the Harvard Center for Comparative Medicine or obtained from Jackson Laboratories. Mice were housed in groups of up to five animals and maintained on a 12hr light/dark cycle. Transgenic alleles were heterozygous for each transgene for all experimental animals. Descriptions of allele combinations for all experiments can be found in **Supp. Table 1**.

Spiral ganglion neurons (SGNs) and their central axons, i.e. auditory nerve fibers, were targeted using either *Foxg1*^tm1.1(Cre)Ddmo^ (*Foxg1*^Cre^)^114^ or *Foxg1*^Flp^, both of which drive robust reporter expression in neurons in the auditory and vestibular ganglion^115,116^ and the neocortex^117,118^, but not in brainstem or midbrain neurons. *Foxg1*^Flp^ mice were generated by crossing the *Foxg1*^tm1.1Fsh^ mouse line^119^ with the Tg(EIIa-Cre)C5379Lmgd mouse line^120^, then backcrossing to isolate the Flp transgene and remove the Cre transgene.

Inhibitory inputs to octopus cells were targeted with *Slc6a5*^tm1.1(Cre)Ksak^ mice (*Glyt2*^Cre^)^121^.

Octopus cells were sparsely labeled with the Tg(Thy1-YFP)HJrs (*Thy1*) mouse line^122^. This line labels ∼0-15 octopus cells amongst other neurons throughout the brain.

Ib/c SGNs were targeted using the *Ntng1*^em1(Cre)Kfra^ (*Ntng1*^Cre^) mouse line, which drives expression in neurons throughout the nervous system (**Supp. Fig. 1F**) and disrupts expression of the endogenous allele^54^. Auditory brainstem responses in adult *Ntng1*^Cre/+^ mice are normal. Ic SGNs were sparsely targeted with the *Myo15a*^tm1.1(Cre)Ugds^ (*Myo15*^iCre^) mouse line^123^.

Fluorescent reporters included *Gt(ROSA)26Sor*^tm14(CAG-tdTomato)^ (Ai14, tdT)^124^, *Gt(Rosa)26Sor*^tm34.1(CAG-Syp/tdTomato)^ (Ai34, syp/tdT), and *Gt(Rosa)26Sor*^tm1.2(CAG-EGFP)Fsh^ (RCE:FRT, EYFP)^125^. We also used *Gt(Rosa)26Sor*^tm32(CAG-COP4*H134R.EYFP)^ (Ai32, ChR2)^126^ to drive synaptic activity in *in vitro* slice experiments.

### Histology and Reconstructions

For immunohistochemical labeling, mice were deeply anesthetized with isoflurane and transcardially perfused with 15mL of 4% paraformaldehyde (PFA) in 0.1M phosphate-buffered saline (PBS) using a peristaltic pump (Gilson). Whole skulls containing brain and cochlea were immediately transferred to 20mL of 4% PFA and post-fixed overnight at 4°C. Fixed brains and cochlea were removed from the skulls and washed with 0.1M PBS.

Brains were collected from mice of both sexes, aged 28-38 days, and embedded in gelatin-albumin hardened with 5% glutaraldehyde and 37% PFA^127^. Sections were cut at 35, 65, or 100μm with a vibrating microtome (Leica VT1000S) and free-floating tissue was collected in 0.1M PBS. For sections less than 65μm, tissue was permeabilized and nonspecific staining was blocked in a solution of 0.2% Triton X-100 and 5% normal donkey serum (NDS, RRID: AB_2337258) in 0.1M PBS for 1 hour. After blocking, tissue was treated with primary antibody in a solution containing 0.2% Triton X-100 and 5% NDS in PBS for 1-2 nights at room temperature. Primary antibodies used were: chicken anti-GFP (1:1000, RRID:AB_10000240), rabbit anti-RFP (1:1000, RRID:AB_2209751), goat anti-calretinin (1:1000, RRID:AB_1000034), and guinea pig anti-VGLUT1 (1:500, RRID:AB_887878). Sections were washed in 0.1M PBS then incubated in a secondary antibody solution (1:1000) containing 0.2% Triton X-100 and 5% NDS for 2-3hrs at room temperature. Tissue sections were mounted on charged slides and coverslipped (Vectashield Hardset Antifade Mounting Medium with DAPI), and imaged using a Zeiss Observer.Z1 confocal microscope.

For 100μm sections, tissue was washed in CUBIC-1A solution^128^ for 1hr for strong permeabilization and delipidization^128,129^. Tissue was then further permeabilized and nonspecific staining was blocked in a solution of 0.2% Triton X-100 and 5% NDS in 0.1M PBS for 1hr. After blocking, tissue was treated with primary antibody in a solution containing 0.2% Triton X-100 and 5% NDS in PBS for 4 nights at 37°C. Primary antibodies used were: chicken anti-GFP (1:1000, RRID:AB_10000240), and rabbit anti-RFP (1:1000, RRID:AB_2209751). Sections were then incubated in a secondary antibody solution (1:400) containing 0.2% Triton X-100 and 5% NDS for 4 nights at 37°C. Tissue sections were pre-incubated in CUBIC2 solution, then temporarily mounted on uncharged slides with CUBIC2 solution for immediate imaging using a Zeiss Observer.Z1 confocal microscope.

Octopus cells and synaptic puncta were reconstructed in Imaris (Oxford Instruments). YFP signal from the target octopus cell was used to generate a surface reconstruction and mask syp/tdT signal. Dendrites were reconstructed using the masked YFP signal and separated into 10μm increments. Masked syp/tdT puncta were marked and localized to a 10μm increment of the dendritic tree. Synapse counts, dendrite metrics, and masked channels were exported to Excel (Microsoft) for further analysis.

Cochlea were collected from mice of both sexes, aged 28-42 days. The bony labyrinth of the inner ear was decalcified in 0.5M ethylenediamine tetraacetic acid (EDTA) for 3 nights at 4°C and embedded in gelatin-albumin hardened with 5% glutaraldehyde and 37% PFA. Sections were cut at 65μm with a vibrating microtome (Leica VT1000S) and freefloating tissue was collected in 0.1M PBS. Sections were washed in CUBIC-1A solution for 1hr for strong permeabilization and delipidization. Tissue was further permeabilized and nonspecific staining was blocked in a solution of 0.2% Triton X-100 and 5% NDS in 0.1M PBS for 1hr. After blocking, tissue was treated with primary antibody in a solution containing 0.2% Triton X-100 and 5% NDS in PBS for 2 nights at room temperature. Primary antibodies used were: chicken anti-GFP (1:1000, RRID:AB_10000240), rabbit anti-RFP (1:1000, RRID:AB_2209751), goat anti-calretinin (1:1000, RRID:AB_1000034). Sections were then incubated in a secondary antibody solution (1:500) containing 0.2% Triton X-100 and 5% normal goat serum for 2-3hrs at room temperature. Tissue sections were mounted on charged slides, coverslipped (Vectashield Hardset Antifade Mounting Medium with DAPI), and imaged using a Zeiss Observer.Z1 confocal microscope.

### Acute Slice Electrophysiology

Data were obtained from mice of both sexes, aged 24-47 days. Mice were deeply anesthetized with isoflurane and perfused transcardially with 3mL of 35°C artificial cerebral spinal fluid (ACSF; 125mM NaCl, 25mM glucose, 25mM NaHCO_3_, 2.5mM KCl, 1.25mM NaH_2_PO_4_, 1.4Mm CaCl_2_, and 1.6mM MgSO_4_, pH adjusted to 7.45 with NaOH). For high calcium concentration experiments presented in **Fig. 3C**, ACSF contained 125mM NaCl, 25mM glucose, 25mM NaHCO_3_, 2.5mM KCl, 1.25mM NaH_2_PO_4_, 2.4mM CaCl_2_, and 1.3mM MgSO_4_. Mice were rapidly decapitated and the brain was removed and immediately submerged in ACSF. Brains were bisected and 250μm slices were prepared in the sagittal plane with a vibrating microtome (Leica VT1200S; Leica Systems). Prepared slices were incubated for 30min at 35°C, then allowed to recover at room temperature for at least 30min. ACSF was continuously bubbled with 95% O_2_/5% CO_2_.

Whole-cell recordings were conducted at 35°C using a Multiclamp 700B (Molecular Devices) in current-clamp mode with experimenter adjusted and maintained bridge balance and capacitance compensation. Data were filtered at 12kHz, digitized at 83–100kHz, and acquired using pClamp9 (Molecular Devices). Neurons were visualized using infrared Dodt gradient contrast (Zeiss Examiner.D1; Zeiss Axiocam 305 mono). Glass recording electrodes (3–7MΩ) were wrapped in parafilm to reduce capacitance and filled with an intracellular solution containing 115mM Kgluconate, 4.42mM KCl, 0.5mM EGTA, 10mM HEPES, 10mM Na_2_Phosphocreatine, 4mM MgATP, 0.3mM NaGTP, and 0.1% biocytin, osmolality adjusted to 300mmol/kg with sucrose, pH adjusted to 7.30 with KOH. All membrane potentials are corrected for a -11mV junction potential.

For optogenetic activation, full-field 475nm blue light was presented through a 20x immersion objective (Zeiss Examiner.D1). Onset, duration, and intensity of light was controlled by a Colibri5 LED Light Source (Zeiss). Light intensity at the focal plane ranged between 1.9 and 4.1mW/mm^2^, corresponding to 6% and 10% intensity on the Colibri5 system. For electrical stimulation, glass stimulating electrodes were placed in the auditory nerve root and 20µs current pulses were generated with a DS3 current stimulator (Digitimer). Light or electrical stimulation intensity was adjusted during the experiment to evoke subthreshold EPSPs, not spikes. During analysis, EPSPs and spikes were distinguished by screening all events with a phase plot analysis. Only stimulation events that evoked EPSPs unambiguously were included for analysis. When presenting electrical and light stimulation together, a series of stimulation pairings with shuffled onset timings was presented to account for cell to cell variability in EPSP and IPSP timings. Data presented is for the stimulation pairings that evoked a maximal shift in EPSP timings.

### Analysis and Statistical Tests

Cell counts and habenula measurements were performed in ImageJ/FIJI software (National Institutes of Health). Electrophysiology data were analyzed using custom scripts and NeuroMatic analysis routines^130^ in Igor Pro (Wavemetrics).

For data with equal variance (Levene’s test), one-way ANOVAs with Tukey’s HSD post hoc test were used where appropriate to determine statistical significance. For data with non-homogenous variances, oneway ANOVAs with a Welch F test were used with a Tukey’s HSD post hoc test. Errors and error bars report standard deviation (SD) or standard error of the mean (SEM) as noted in figure legends and throughout the text.

### Computational Modelling

Computer simulations were performed using the NEURON 8.2 simulation environment^131^, with an integration time constant of 25µs. The morphology of the octopus neuron was obtained from McGinley et al., 2012^30^. The active and passive properties of the model were optimized to match the experimental recordings. We set the passive parameters as follows: internal or axial resistance (R_i_ or R_a_) to 150Ω.cm, membrane resistance (R_m_) to 5KΩ.cm^2^, capacitance (C_m_) to 0.9µF/Cm^2^ and resting membrane potential (V_m_) to -65mV. Ion-channel kinetics, maximum conductance densities, Q10 (3), and temperature (22°C) were obtained from and matched to Manis and Campagnola, 2018^71^ and the maximal conductances were adjusted using a scaling factor (scl) to align qualitatively with experimental data (**Supp. Fig. 4A-B**): fast 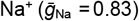, fast transient 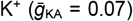, high threshold 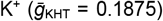, low-voltage activated 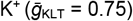, hyperpolarization-activated cyclic nucleotide-gated 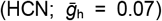 and leak 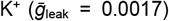. All conductances were uniformly distributed across dendrites and soma, except for HCN conductance, which was only present in dendrites. A baseline scaling factor of 1 was applied under control conditions and 0 under Kv+HCN block conditions. AMPA conductance (*G*_AMPA_) was set to 5nS to align with experimental data (**Fig. 5F**), and glycine conductance (*G*_Gly_) was set to 1nS to match experimental observations (**Fig. 4E**). Reversal potentials for HCN, Na^+^, and K^+^ respectively were (in mV), E_h_= -38, E_Na_= 50 and E_K_= -70. Excitatory AMPA synaptic conductance and inhibitory glycine synaptic conductance were introduced in the proximal and distal dendrites to test the impact of dendritic inhibition on the EPSP height and peak time. The magnitudes of synaptic conductances were tuned to fall within the range seen during experimental data collection. The rise and decay time of AMPA and glycine conductances were tuned to 0.3ms and 3ms respectively, to match experimental data. The reversal potential of AMPA and glycine conductance was set to 0mV and -80mV respectively.

---

**Supplementary Table 1.**
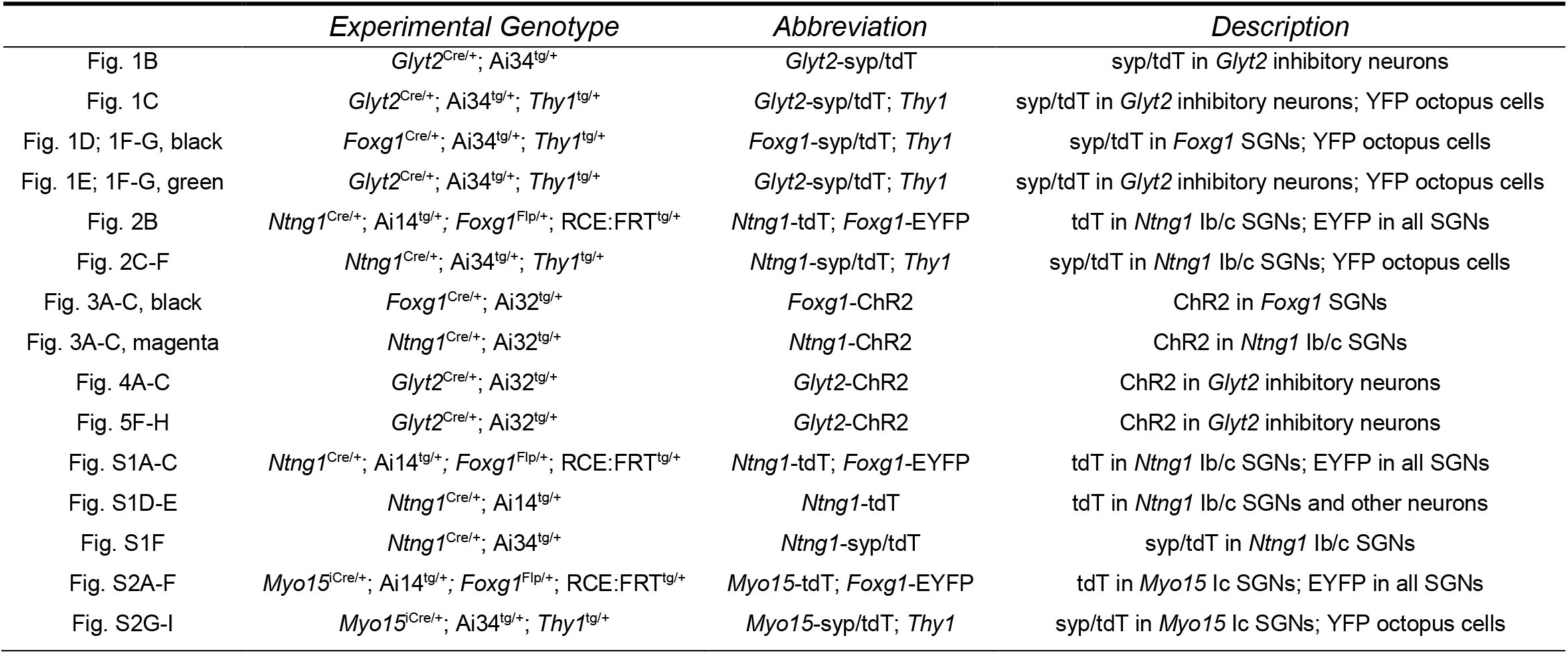
Summary and description of experimental genotypes presented in figures.

**Supplementary Figure 1.**
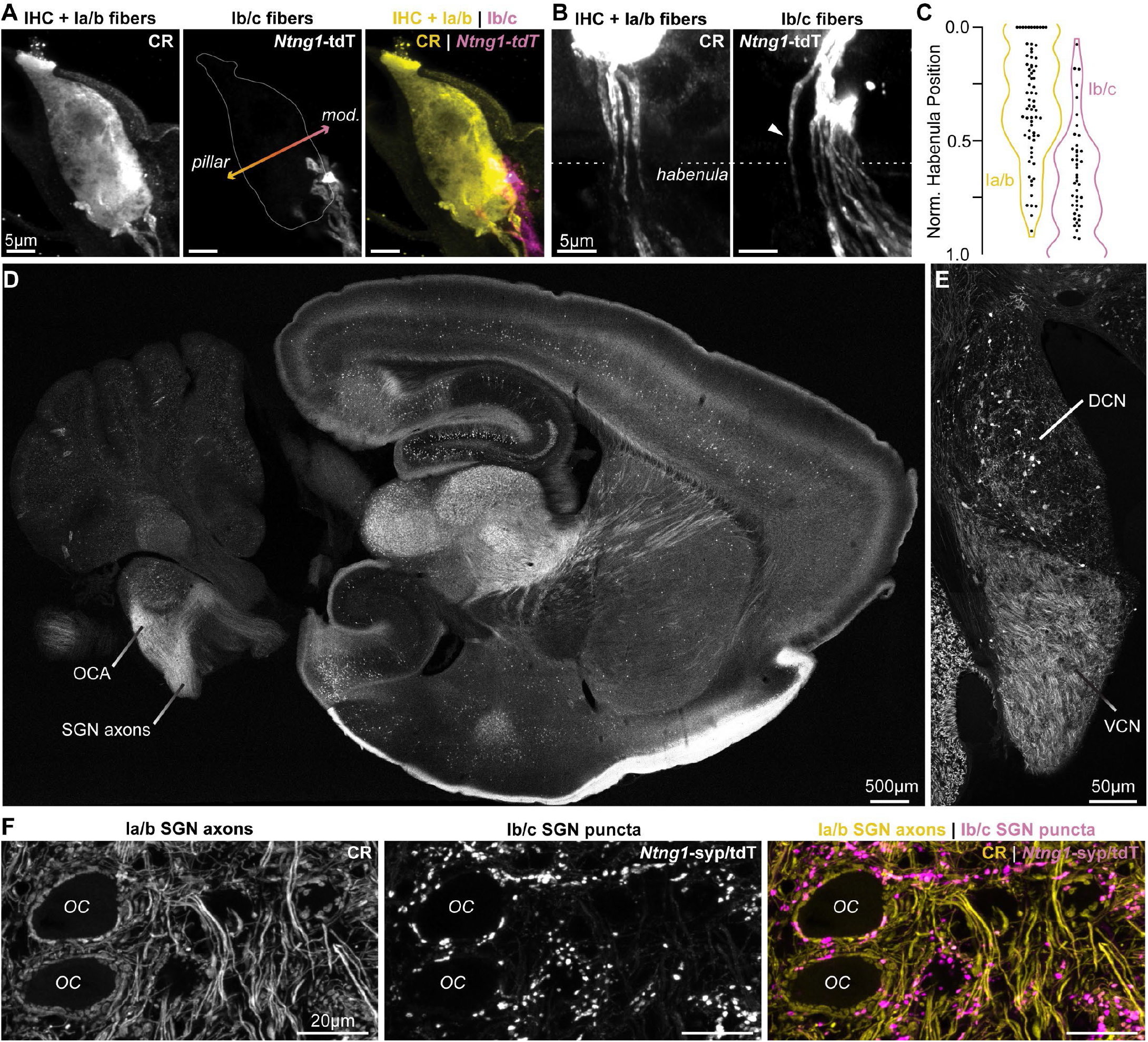
*Ntng1*^Cre^ has high specificity for Ib/c SGNs. **(A)** Calretinin immunopositive (CR+) Ia/b spiral ganglion neuron (SGN) peripheral fibers (CR, yellow) preferentially innervate the pillar side of inner hair cells (IHCs). Ib/c SGN peripheral fibers with *Ntng1*^Cre^-mediated expression of tdTomato (*Ntng1*-tdT, magenta) preferentially innervate the modiolar side of IHCs. IHCs also immunolabel for CR. **(B)** CR+ Ia/b SGN fibers (yellow) pass through the pillar side of the habenula while *Ntng1*-tdT+ Ib/c SGN fibers (magenta) pass through the modiolar side. Arrowhead highlights a *Ntng1*-tdT fiber passing through the pillar side of the habenula but ultimately terminating on the modiolar side of the IHC. **(C)** Normalized position of CR+ Ia/b (yellow) and *Ntng1*-tdT+ Ib/c (magenta) fibers in the habenula (*n* = 124 fibers; 5 mice). **(D)** In the central nervous system, *Ntng1*-tdT is present throughout the brain, but is restricted to SGN axons in the ventral cochlear nucleus where the octopus cell area (OCA) is found. **(E)** A coronal section of the cochlear nucleus highlights *Ntng1*-tdT SGN axons in the ventral cochlear nucleus (VCN) and scattered *Ntng1*-tdT somas in the dorsal cochlear nucleus (DCN). **(F)** In the octopus cell area, CR immunolabel is present in Ia/b SGN axons and puncta. As in the ganglion, CR co-labels with some Ib/c puncta (*Ntng1*-syp/tdT).

**Supplementary Figure 2.**
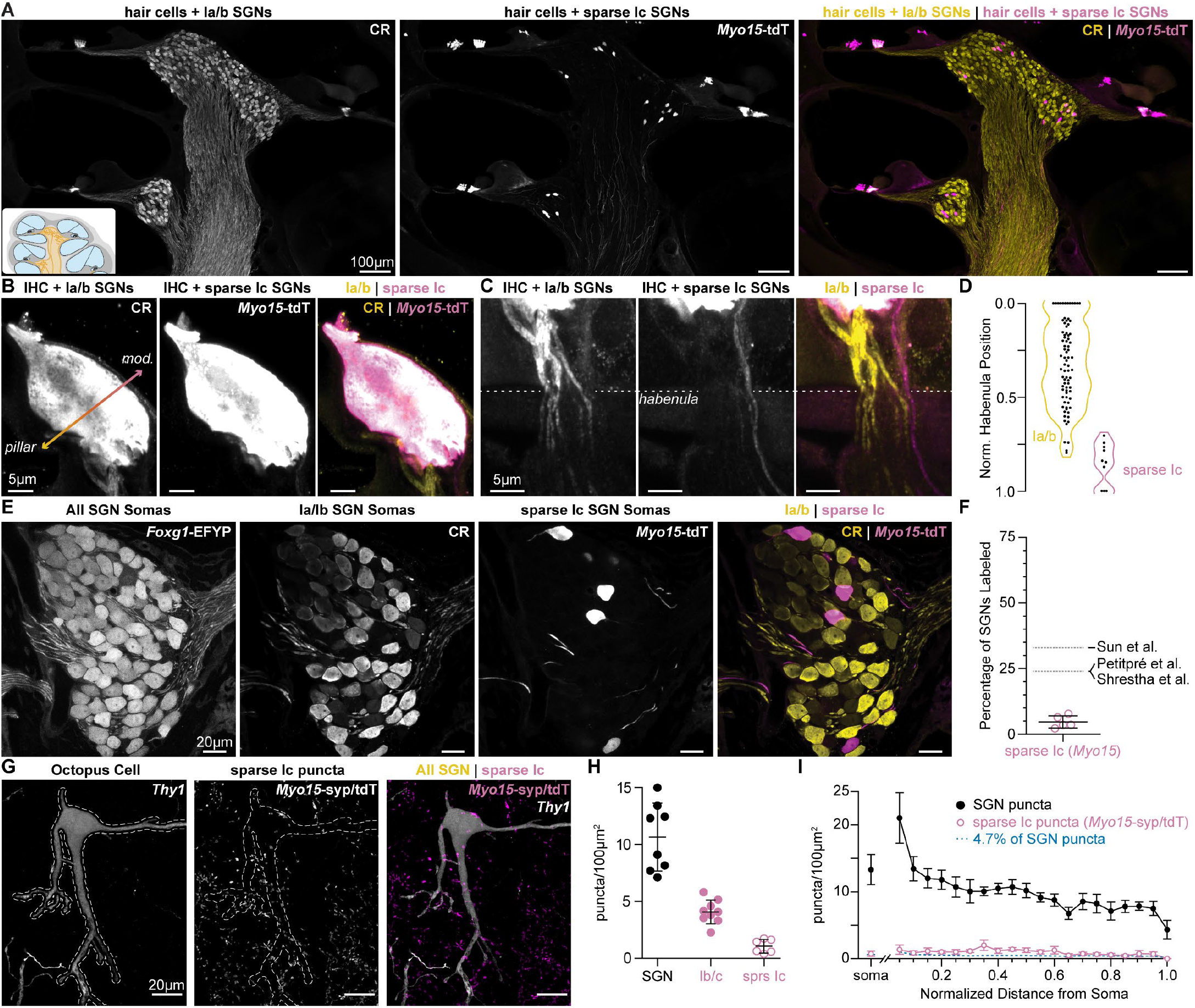
*Myo15*^iCre^ sparsely labels Ic SGNs. **(A)** Cochlear sections with calretinin (CR) immunolabeling of hair cells and type Ia/b spiral ganglion neurons (SGNs) and *Myo15*^iCre^-mediated expression of tdTomato (*Myo15*-tdT) in hair cells and some type Ic SGNs. **(B)** CR+ Ia/b SGN fibers (yellow) preferentially innervate the pillar side of inner hair cells (IHCs). Sparse Ic SGN fibers with *Myo15*-tdT (magenta) preferentially innervate the modiolar side of IHCs. IHCs label with both tdT and CR. **(C)** CR+ Ia/b SGN fibers (yellow) pass through the pillar side of the habenula while sparsely labeled *Myo15*-tdT+ Ic SGN fibers (magenta) pass through the modiolar side. **(D)** Normalized position of CR+ Ia/b (yellow) and *Myo15*-tdT+ Ic (magenta) SGN fibers along the pillar to modiolar axis of the habenula (*n* = 90 fibers; 4 mice). **(E)** 65µm cochlear section containing SGN somas. SGNs have variable levels of calretinin (CR) immunolabeling corresponding to the three molecular subtypes. Ia/b SGN somas label with high and medium levels of CR, respectively. Ic somas label with very low levels of CR. *Myo15*-tdT is sparsely found in Ic SGNs. All SGN somas are labeled with *Foxg1*^Flp^-mediated expression of EYFP (*Foxg1*-EYFP). **(F)** tdT+CRSGNs make up 4.7 ± 2.3% of the SGN population (*n* = 2150 neurons, 5 mice), indicating sparse reporter expression. Data are presented as mean ± SD; individual data points signify percent coverage per animal. Dotted lines are estimated percentages for type Ic SGNs from 1: Petitpré et al., 2018, 2: Shrestha et al., 2018, and 3: Sun et al., 2018. **(G)** A sparsely labeled *Thy1* octopus cell has few *Myo15*-syp/tdT puncta near its soma and dendrites. **(H)** Density of all SGN (black: data from Fig. 1D), Ib/c SGN (magenta: data from Fig. 2D), and sparse Ic SGN inputs (open magenta circles: 1.1 ± 0.6, *n* = 6 cells, 3 mice). Data are presented as mean ± SD. Markers represent the total puncta density computed per reconstructed octopus cell. **(I)** Puncta density per 100µm^2^ of soma surface area (all SGN, black: data from Fig. 1E; sparse I/c inputs, magenta open circles: 0.7 ± 0.4, *n* = 6 cells, 3 mice) and density along the length of the dendritic tree, relative to the soma. Data are presented as mean ± SEM.

**Supplementary Figure 3.**
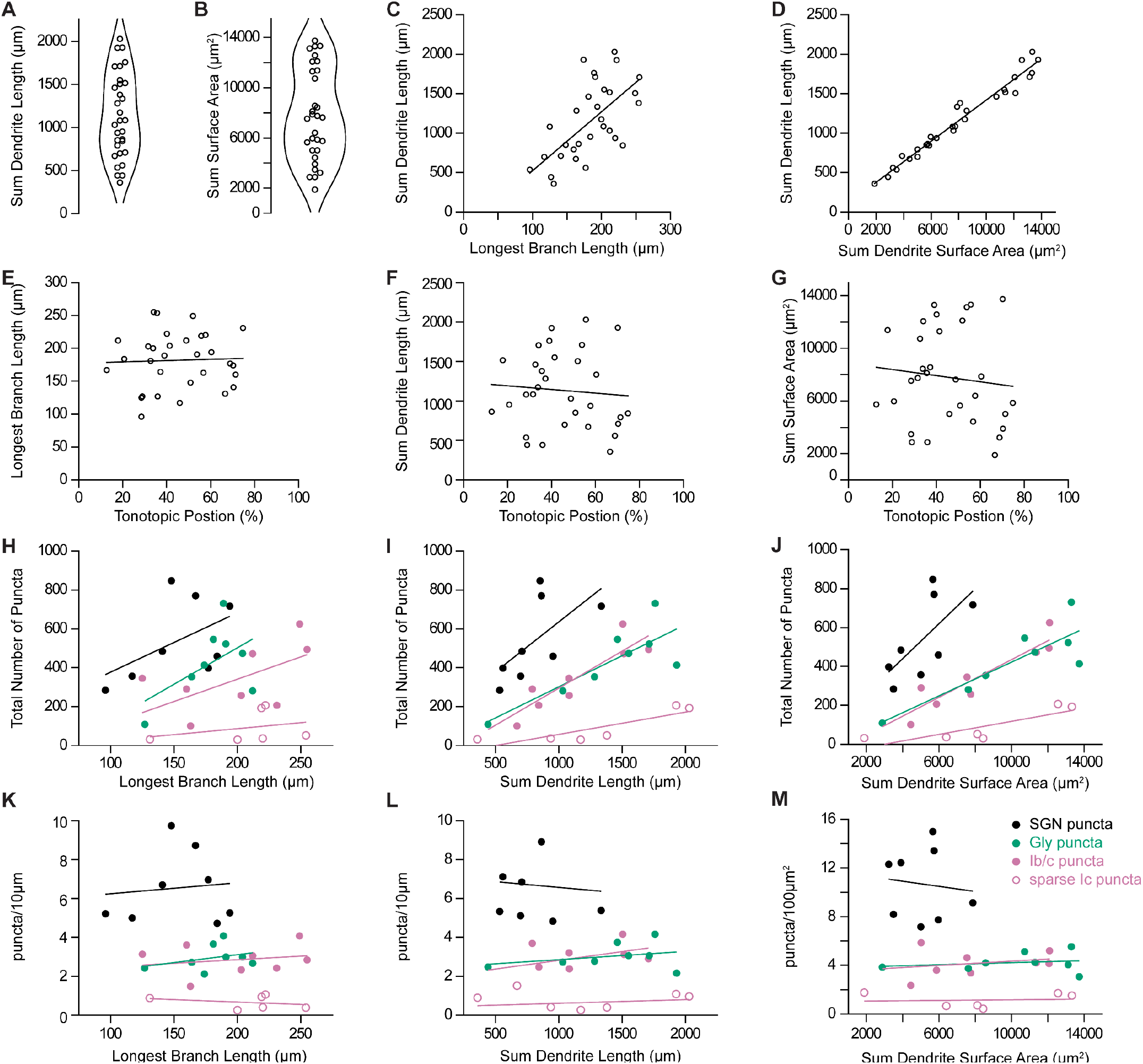
Dendritic and synaptic reconstructions of octopus cells. **(A)** Total length of reconstructed dendritic arbors for 31 octopus cells. **(B)** Total surface area of reconstructed dendritic arbors for 31 octopus cells. **(C)** Octopus cell reconstructions were normalized to the longest reconstructed dendrite. Total length of reconstructed dendrites correlated with the longest branch per neuron. **(D)** Total length of reconstructed dendrites correlated with total dendritic surface area. **(E-G)** Longest branch length, total dendrite length, and total surface area compared to estimated position of the octopus cell soma in the tonotopic organization of the octopus cell area. **(H-J)** Total number of reconstructed spiral ganglion neuron (SGN) puncta (*Foxg1*-syp/tdT, black), inhibitory puncta (*Glyt2*-syp/tdT, green), Ib/c SGN puncta (*Ntng1*-syp/tdT, magenta), and sparse Ic SGN puncta (*Myo15*-syp/tdT, open magenta circles) compared to the longest branch length, total dendrite length, and total dendrite surface area. **(K-M)** Density of reconstructed SGN puncta (*Foxg1*-syp/tdT, black), inhibitory puncta (*Glyt2*-syp/tdT, green), Ib/c SGN puncta (*Ntng1*-syp/tdT, magenta), and sparse Ic SGN puncta (*Myo15*-syp/tdT, open magenta circles) compared to longest branch length, total dendrite length, and total dendrite surface area.

**Supplementary Figure 4.**
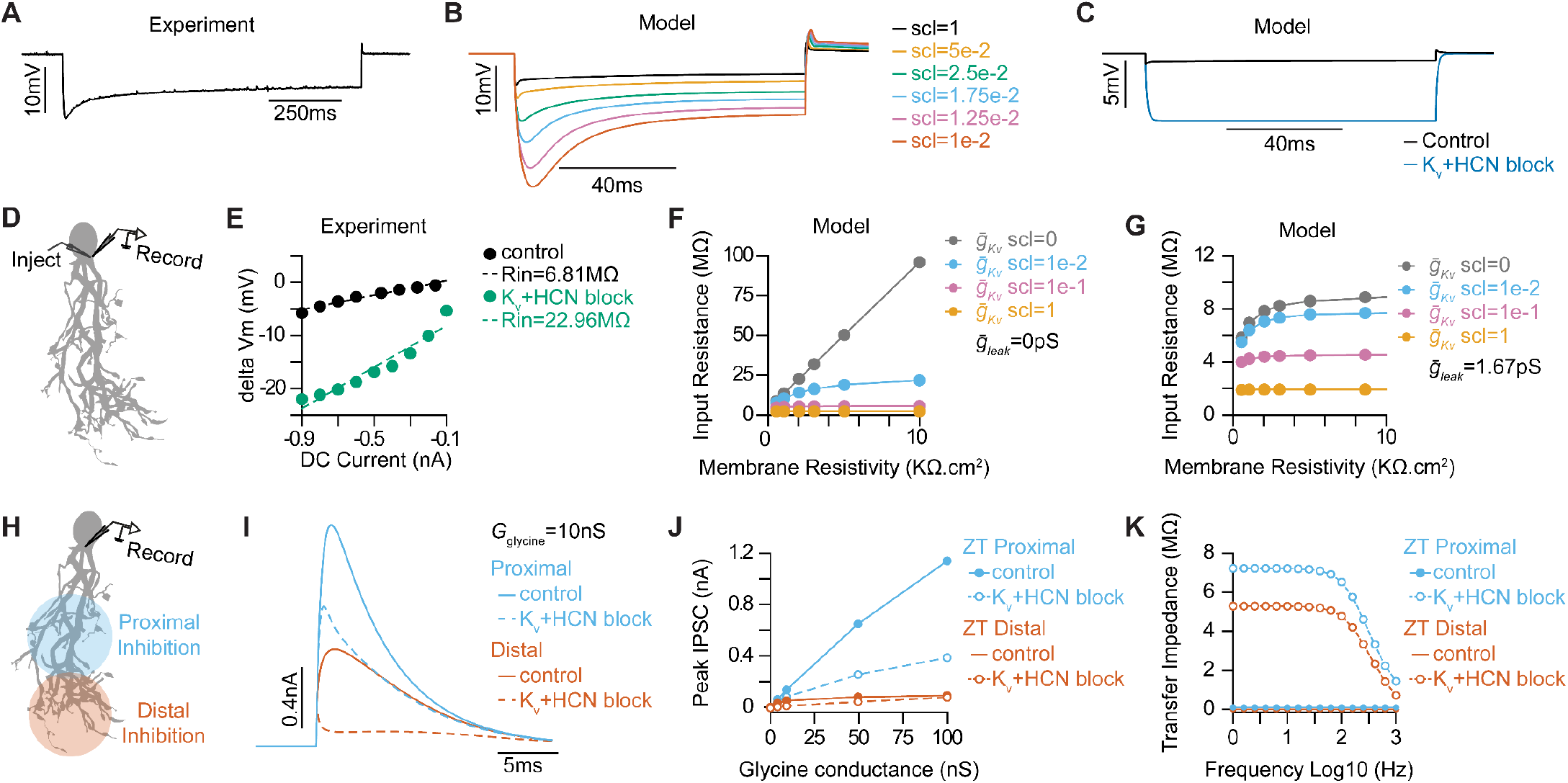
Optimizing active and passive properties of an octopus cell model. **(A)** Subthreshold somatic voltage response to a -800pA hyperpolarizing current injection in an *in vitro* whole-cell current clamp recording of an octopus cell. **(B)** Somatic voltage responses from a morphologically realistic octopus cell model to a -800pA current injection for various scaling factors (scl) of maximal conductances of 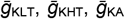 and 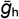.**(C)** Comparison of somatic hyperpolarizations to a -800pA current injection in the model reproducing experimental data during control (black) and Kv and HCN block conditions (blue). **(D)** Illustration of injection and recording locations for panels *E-G* in a morphologically realistic octopus cell model. **(E)** Current-voltage (IV) relationships from a representative experimental octopus cell recorded in control ACSF and with Kv and HCN block. Dotted lines plot linear fits of the experimental data. The slope of the dotted line estimates Rin. **(F-G)** Impact of leak conductance 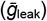 and scaling factor (scl) on passive properties of the model with 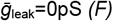 and 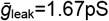 *(G)* for various scaling factor values (indicated in different colors). **(H)** Illustration of injection and recording locations for panels *I-K* in a morphologically realistic octopus cell model. **(I)** Somatically recorded inhibitory post synaptic currents (IPSCs) recorded from glycinergic synapses in proximal (blue) and distal (orange) stimulation during control (solid) and Kv and HCN block conditions (dotted). **(J)** Peak IPSC magnitude as function of glycine conductance in proximal (blue) and distal (orange) stimulation during control (solid) and Kv and HCN block conditions (dotted). **(K)** Transfer impedance as function of frequency from proximal dendrites to soma (blue) and distal dendrites to soma (orange) during control (solid) and Kv and HCN block conditions (dotted).

**Supplementary Figure 5.**
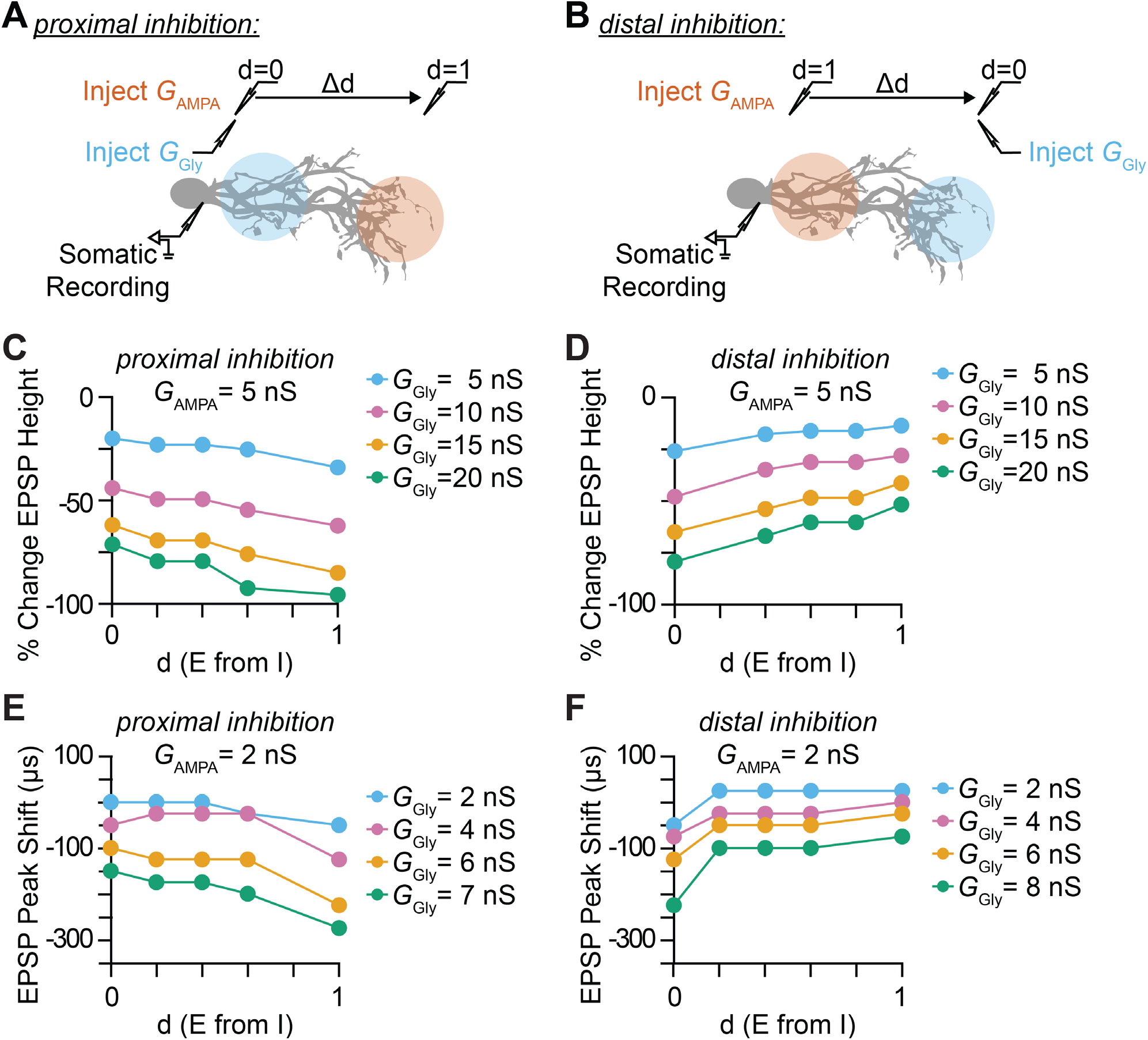
Impact of inhibitory synaptic location and distance between excitatory and inhibitory synapse on somatic EPSP amplitude and timing. **(A-B)** Illustration of injection and recording locations for proximal *(A)* and distal *(B)* inhibition paradigms and the normalized relative distance (d) between excitatory synapses. The impact that the location of dendritic inhibition has on somatic EPSPs is primarily determined by the local potential change by an EPSP and the attenuation or the length constant (λ) of the IPSP towards the excitatory synaptic location. The exponential decay of membrane voltage is asymmetric, with lower λ for the open end and higher λ for sealed end propagation. Distal parts of the dendrites have higher local input resistance and lower attenuation of IPSP due to the sealed end. **(C-D)** Percentage change in somatic EPSP height with dendritic glycinergic inhibition as function of normalized distance between excitatory and inhibitory synapses in proximal *(C)* and distal *(D)* inhibition for various E/I ratio with excitatory AMPA conductance (*G*_AMPA_) set at 5nS. Average shown in black with SEM in shaded region. **(E-F)** Somatic EPSP peak time shift with dendritic glycinergic inhibition as function of normalized distance between excitatory and inhibitory synapses in proximal *(E)* and distal *(F)* inhibition for various E/I ratio with excitatory AMPA conductance (*G*_AMPA_) set at 2nS. Average shown in black with SEM in shaded region.

